# Structure of a secreted lancelet carotenoid cleavage dioxygenase (CCD), from an ancestral CCD clade

**DOI:** 10.64898/2026.10.29.762971

**Authors:** Sheetal Uppal, Gleysin Cabrera Herrera, Liya Muslinkina, Gowthaman Govindarajan, Yan Li, David N. Garboczi, Igor B. Rogozin, Eugenia Poliakov, Apostolos G. Gittis, T. Michael Redmond

## Abstract

Carotenoid cleavage dioxygenases (CCDs) are a superfamily of non-heme iron-containing enzymes ubiquitous across all kingdoms of life. Previously, we identified a new subfamily of CCDs termed beta-carotene oxygenase-like (BCOL) proteins, a chordate expansion of an ancestral sister clade of metazoan CCDs, which are widely distributed in marine invertebrates, including mollusks and lancelets. Here we present the three-dimensional structure of a representative BCOL protein from the lancelet *Branchiostoma fforidae* (BCOLg) at 2.4 Å resolution and its functional properties. BCOLg symmetrically cleaves β-carotene into all-*trans* retinal. RNAseq of wild-caught animals detected six of the seven annotated BCOL transcripts in *Branchiostoma* gut tissue. We expressed and crystallized recombinant BCOLg. BCOLg is heavily glycosylated, with four N-linked glycosylation sites, and adopts the seven-bladed propeller fold characteristic of the superfamily, with one disulfide bond contributing to structural stability. BCOLg and several other BCOL proteins contain Asp substitutions at two of three Glu positions found in other CCDs, stabilizing three of four His residues involved in Fe^2+^ coordination. Notably, BCOLg lacks the canonical -PDPCK-motif characteristic of other metazoan CCDs.

**Summary:** Reporting the structure of lancelet beta-carotene oxygenase-like BCOLg protein at 2.4 Å, an unusual carotenoid cleavage dioxygenase (CCD) superfamily member. BCOLg is a heavily glycosylated protein with a distinctive iron-coordination environment and lacking the canonical metazoan -PDPCK-motif.

## Introduction

The carotenoid cleavage dioxygenases (CCDs) are a superfamily of non-heme iron-containing oxygenases present in all kingdoms of life ^1^. All CCDs, except for the retinal pigment epithelium (RPE) retinoid isomerase RPE65, function in the oxidative cleavage of double bonds ^2^. Closely related stilbene cleavage oxygenases (SCOs) in bacteria, such as lignostilbene dioxygenase (LSD), catalyze the cleavage of the double bonds of stilbenes (e.g., lignostilbene and resveratrol). The CCD superfamily has evolved to oxidatively cleave double bonds of various terpenoids (isoprenoids) and rarely to isomerize *trans* to *cis* in conjugated polyene systems ^1,3–5^. Plant CCDs are involved in a diversity of carotenoid secondary metabolism, such as in the first committed step of abscisic acid biosynthesis, an important phytohormone, formation of strigolactone, another phytohormone, and generation of various plant scent compounds, etc. Metazoan CCDs are similarly diverse in their functions and substrates and are responsible for asymmetric xanthophyll cleavage (BCO2), symmetric provitamin A carotenoid cleavage to produce vitamin A (BCO1), and additionally, for isomerization of all-*trans* retinyl ester in the vertebrate visual cycle (RPE65) ^2,6,7^. Remarkably, insect NinaB can both cleave and isomerize its xanthophyll substrates^4^. Sequence identity between various animal homologs is around 40%. Yet, the precise functions/substrates of many ancestral metazoan CCDs are still undetermined^2^.

In 2017, we described a clade of BCO2-like oxygenases ^1^. Their sequence identity to classic animal CCDs falls in the <30% range. The BCOL ancestral clade within the animal CCD superfamily does not have the membrane -PDPCK-anchoring motif that is highly conserved in other animal CCDs. In RPE65 and BCO2 the cysteine of -PDPCK-is palmitoylated for association with the appropriate intracellular membrane^5^. BCOLs were predicted by modeling to have the same general seven-bladed propeller fold with coordination of non-heme iron by four histidines^1^. Purification and crystallization of recombinant animal CCDs remain challenging due to the enzymes’ intrinsic hydrophobicity and tendency to associate with membranes or lipid aggregates^4^. To date, crystal structures have been determined only for recombinant CCDs from *C. elegans* and insects (NinaB)^4,8^. On the other hand, RPE65 from the native bovine RPE has been crystallized^9^, but this has limited application to study human pathological RPE65 mutations. Our original analysis suggested that lancelet BCOLg is a secreted glycosylated enzyme, and we found in a bacterial assay that it can cleave β-carotene. As for many secreted glycosylated enzymes, such as many proteases, BCOLg is predicted to be highly soluble, and this property gave us an advantage in the purification and crystallization of the recombinant protein from mammalian cell culture. Here we describe the crystal structure of chordate BCOLg, that we identified by RNAseq as being expressed in lancelet gut, at 2.4 Å resolution. The protein adopts the predicted seven-bladed β-propeller fold typical of CCDs and is heavily glycosylated, featuring four N-linked glycosylation sites with resolved N-acetylglucosamine (NAG) moieties. While it coordinates non-heme ferrous ion with four histidines, as other CCDs, it uses a novel arrangement of second-shell residues, compared to other CCDs.

## Methods

### Construction of expression plasmids for BCOLg

DNA sequence encoding the wildtype lancelet BCOLg (GenBank code XP_002601288.1), including its natural leader peptide, was cloned into a pcDNA™3.1/V5-His-TOPO® expression vector in the TA cloning site with a V5/His tag at the C-terminus. An updated sequence with 2% difference (XP_035666855.1) has two additional predicted N-glycosylation sites.

#### RNASeq analysis of Lancelet gut

The live animals (*Branchiostoma fforidae*) were received from Gulf Specimen Marine Labs (Panacea, FL) and dissected under light microscope. Dissected tissues were immediately frozen on dry ice. The tissue was homogenized in homogenization buffer and total RNA was isolated according to standard Maxwell Simply RNA tissue kit protocol (Promega Corp.). Three individual samples with RIN 9.9-10 were used for directional mRNA library preparation (poly A enrichment), sequenced by NovaSeq X Plus Series (PE150; ∼100 M reads/30 G raw data per sample), and analyzed with RNAseq GeneCount analysis package from Novogene (Sacramento CA). Data were visualized in IGV 2.16.2 software.

### Protein expression and purification of BCOLg protein from Expi233 and GnTI-cells

The lancelet BCOLg was expressed using the Expi293™ Expression System (ThermoFisher Scientific) according to the manufacturer’s protocol, scaled to a 30 ml working volume. Briefly, Expi293F™ or Expi293F GnTI-cells were maintained in Expi293™ Expression Medium in a humidified incubator at 37 °C with 8% CO_2_, with continuous orbital shaking (125 rpm). Prior to transfection, cell density was adjusted to 3×10^6^ viable cells/ml. Transfection complexes were prepared by diluting 30 µg of plasmid DNA into 1.5 ml of Opti-Plex™ Complexation Buffer and 80 µL of ExpiFectamine™ 293 Reagent into 1.5 ml of the same buffer. After a 5-min incubation, the diluted reagent was added to the DNA, and the mixture was incubated at room temperature for 30-40 min to allow complex formation. The complexes were then added dropwise into the cell culture. At 16-18 h post-transfection, the culture was supplemented with 150 µl of ExpiFectamine™ 293 Transfection Enhancer 1 and 1.5 ml of Enhancer 2. The secreted BCOLg was harvested at three days post-transfection by centrifugation at 1,000 x g for 10 minutes, followed by sequential clarification of the supernatant through 0.45 μm and 0.22 μm filters. Protein expression was verified by immunoblotting and the supernatants were stored at −80 °C.

All purification steps were carried out at 4 °C. Thawed culture supernatant was concentrated using Amicon Ultra centrifugal filters with 30 kDa cutoff. The concentrate was desalted using a 5 ml HiTrap Desalting column (Cytiva) attached to an ÄKTA purifier system (Cytiva). The column was equilibrated with 20 mM Tris-HCl pH 7.4, 150 mM NaCl, at a flow rate of 2.5 ml/min. Eluted fractions were subjected to SDS-PAGE analysis and fractions containing BCOLg were pooled to ∼20 ml total and concentrated. The sample was then centrifuged at 10,000 × g for 1 min before loading onto a HiLoad 16/60 Superdex 200 column (Cytiva) equilibrated with 20 mM Tris-HCl pH 7.4, 150 mM NaCl, 1 mM TCEP, and 4% glycerol. Size exclusion chromatography was performed at a flow rate of 1 ml/min, collecting fractions across 1 column volume (125 ml). Fractions corresponding to the major elution peak were analyzed by SDS-PAGE and the final pooled protein fractions were concentrated to a final concentration of 6.7 mg/ml (Expi293) and 6.4 mg/ml (GnTI-). Protein concentration was determined by either Bradford assay or from A_280_ using the calculated extinction coefficient ε_280_ of 64,290 M^−1^ cm^−1^.

#### Immunoblotting

BCOLg protein samples were mixed with 4× NuPAGE LDS sample buffer containing SDS, heated at 95 °C for 5 min, briefly centrifuged, and separated on 4–12% Bis-Tris NuPAGE gels (Invitrogen, Thermo Fisher Scientific). Proteins were transferred onto nitrocellulose membranes using the iBlot 2 Dry Blotting System (Invitrogen). Membranes were blocked with Intercept Blocking Buffer (LI-COR Biosciences) and incubated overnight at 4 °C with primary antibodies against the 6×His tag (anti-mouse mAb (Cat. No. 2366 1:1000), anti-rabbit mAb (Cat. No. 12698 1:1000) Cell Signaling Technology) and HSP70 (anti-mouse pAb (Cat. No. ADI-SPA-812 1:1000) Enzo Biochem Inc). After washing, membranes were incubated with the appropriate fluorescent secondary antibodies. Protein bands were visualized using the Odyssey M Imaging System (LI-COR Biosciences, Lincoln, NE, USA).

### In vitro activity assay for BCOLg using β-carotene substrate

*In vitro* assays for BCOLg activity were performed based on the method of Lindqvist and Andersson^10^ with some modifications described previously^11^ and the addition of 1% n-decyl-ß-D-maltoside (DMP). Typically, reactions were run in glass vials under reduced light in a volume of 200 μl in 100 mM Tricine-KOH pH 8.0 buffer supplemented with 10 μM FeSO_4_, 5 mM Tris(2-carboxyethyl)phosphine hydrochloride (TCEP), 1% (w/v) n-octyl-β-D-thioglucopyranoside (OTG), 1% glycerol and 1% DMP. The enzyme concentration varied between 80-200 µg/reaction, as determined by BCA protein assay. The concentration of β-carotene in the reaction mixture was 12 µM. The substrate was solubilized in hexane with 50 μl of 4% OTG in a glass vial, and the solvent was evaporated under argon. The assay buffer with substrate/detergent was preincubated at 37 °C for 5 min, enzyme added, and the reaction incubated at 37 °C in a water bath for 45 min. To analyze reaction products, we used a protocol modified from During et al^12^: 50 μl of 37% (v/v) formaldehyde was added to stop the reaction, and the incubation was continued for 10 min at 37 °C. Then, 500 μl of acetonitrile was added, the solution vortexed and put on ice at 4 °C for 5 min. The upper acetonitrile phase was collected and 100 μl injected and analyzed by RP-HPLC.

### HPLC analysis of carotenoids and all-trans retinal

β-carotene, all-*trans* retinal and β-apo-8’-carotenal (internal standard for *in vivo* assays) were separated on a 4.6 × 250 mm 3 μm YMC C30 carotenoid column (Waters, Inc.) at a flow rate of 1 ml/min and simultaneous UV detection at 451 nm and 383 nm (Agilent 1200 HPLC), as described previously with small modifications^11^. Briefly, the initial conditions consisted of 60:40 acetonitrile/0.015 M NH_4_OAc held for 5 min, followed by a linear gradient to 50:50 isopropanol/acetonitrile in 10 min. This was held for an additional 10 min and then followed by a linear gradient to 80:20 isopropanol/acetonitrile in 5 minutes and held for an additional 2 minutes before returning to initial conditions.

β-carotene and all-*trans* retinal were quantified from their peak area using standard curves obtained with 2-220 ng and 0.5-20 ng of material, respectively, in 60/40 acetonitrile:0.015 M ammonium acetate.

### Circular Dichroism (CD) Spectroscopy

To determine the effects of the expression system and buffer composition on the secondary structure of BCOLg, far-UV circular dichroism (CD) spectroscopy was performed using BCOLg expressed in Expi293 and GnTI⁻ cells. For CD measurements, the Expi293-derived BCOLg was analyzed either in the original Tris buffer containing glycerol or after dilution into glycerol-free Tris buffer, whereas the GnTI^−^-derived BCOLg was analyzed in glycerol-free Tris buffer. Protein samples were diluted to a final concentration of 0.1 mg mL⁻¹ (approximately 1.6 μM). Far-UV CD spectra were recorded using a Chirascan Ǫ100 spectropolarimeter (Applied Photophysics) in a 1 mm path-length quartz cuvette over the wavelength range of 190-260 nm with a 1 nm bandwidth at 25 °C. Three consecutive scans were collected for each sample, averaged, and corrected by subtracting the corresponding buffer spectrum, followed by noise reduction.

### Mass Photometry

Mass photometry (MP) measurements were performed as described previously ^13^. Coverslips (24 × 50 mm) were cleaned sequentially with distilled water, ethanol, and isopropanol, followed by drying with a stream of clean nitrogen. The optical quality of each coverslip was verified by imaging a drop of distilled water. Flow chambers were assembled from stacked coverslips and double-sided tape, while silicone gasket wells were prepared by rinsing the gasket with water and placing it on the working side of the coverslip, pressing gently with a clean pipette tip. All buffers were filtered through 0.22 μm syringe filters prior to use. Purified BCOLg protein stock solution was centrifuged at maximum tabletop speed for 10 min to remove aggregates. For MP measurements, 100 μl protein samples were prepared at final concentrations of 20 nM in flow chambers. Video files were processed using the instrument’s software to identify and quantify molecular landing events. Filter parameters were adjusted as needed to optimize event detection and minimize background noise based on the molecular mass range analyzed. Temporal changes in signal intensity and frame appearance were examined to detect artifacts, and spatial or temporal masks were applied to exclude high-noise regions. Contrast values were converted to molecular masses using the instrument’s calibration function, which was routinely validated using an unstained protein standard ladder. Mass distributions were visualized as histograms or kernel density estimates (KDEs), with bin width or bandwidth optimized to resolve distinct components while minimizing noise. Gaussian fitting or KDE peak finding was used to determine the mean molecular masses of each species. All analysis parameters—including sample and buffer details, dilution ratios, final concentrations, chamber type, acquisition time, and modified software settings—were recorded to ensure reproducibility.

### Particle size and polydispersity index

Dynamic light scattering (DLS) measurements were carried out at 25 °C using a Malvern Zetasizer (Malvern Instruments, Ltd., Malvern, U.K.) to determine the mean hydrodynamic diameter and polydispersity index (PDI) of the purified protein samples. The hydrodynamic diameter represents the average particle size, while the PDI reflects the width of the size distribution.

### BCOLg crystallization, X-ray data collection, structure determination and model building

Initial crystallization screening was carried out using commercial kits from Hampton Research (California, USA). Crystals were obtained using the hanging-drop vapor diffusion method at 25 °C. The well solution consisted of 0.2 M sodium acetate trihydrate, 0.1 M Tris-HCl pH 8.5, and 30% (w/v) polyethylene glycol 4000. After 25–30 days of incubation, small needle-like crystal showers appeared. These preliminary conditions were further optimized by testing various additives. Each drop contained 1–2 μl of ∼ 6 mg/ml protein solution mixed with 1 μl of the well solution. Under optimized conditions with 5% ethylene glycol, larger, well-formed crystals appeared within approximately two weeks.

The data were collected on a RIGAKU FRX X-ray source and a Dectris Eiger2R 4M using Cu radiation with ethylene glycol as the cryoprotectant. Crystals diffracted to 2.4A resolution in space group C2. The structure was solved by molecular replacement used a BCOLg model produced by AlphaFold 2.0 (Google CoLab)^14,15^ followed by rigid body fitting. Rebuilding to optimize geometry and stereochemistry yielded R_work_/R_free_ values of 17.5/22.4% (SI Appendix, Table S1). Diffraction data were processed with the XDS^16^ software package^17^. Refinement was carried out with Phenix^18^ and model building was carried out with Coot ^19^. Carbohydrates were modelled into Fo-Fc maps and further refined using Fo-Fc, and 2Fo-Fc maps. The Ramachandran plot as determined by MOLPROBITY ^20^ showed > 89.4% of all residues in favored regions and > 99.5% of all residues in allowed regions. Data collection and refinement statistics are shown in SI Appendix, Table S1. PISA analysis was carried out with Protein interfaces, surfaces and assemblies service PISA at the European Bioinformatics Institute^21^.

### Sequence alignment and analysis of substrate tunnel and comparison to RPEC5 and ACO tunnels

Sequence alignment analysis was performed using ESPript 3.0 server. Tunnel and pocket analyses of the BCOLg crystal structure were performed using MOLEonline. For comparative analysis, the crystal structures of bovine RPE65 (PDB:3FSN) and *Synechocystis* ACO (PDB:2BIW) were used.

### Glycosylation analysis

For N-glycosylation analysis, a band corresponding to BCOLg was excised from the native PAGE gel and subjected to the procedure described by Royle et al. ^22^. Briefly, the BCOLg band was cut into 1 mm³ pieces and placed into 1.5 ml tubes. The gel pieces were destained by incubating three times with 250 μl of 25 mM NH_4_HCO_3_ for 10 minutes at 37 °C, followed by dehydration with 250 μl of pure acetonitrile. For N-glycan release, the gel was rehydrated with 30 μl of PNGase F solution in 25 mM NH_4_HCO_3_ (50 U/μl) and incubated for 16 hours at 37 °C. After the deglycosylation reaction, the gel pieces were washed three times with 200 μl of water for 30 minutes at 37 °C. The supernatants collected after each wash were combined for MALDI-MS analysis.

N-glycans were purified using microcrystalline cellulose solid phase extraction (MCC SPE) before detection in MALDI-MS ^23^. The purified N-glycans were then dried and resuspended in 5 μl of water. For N-glycan detection, 0.5 μl of the N-glycan solution and 0.5 μl of 2,5-dihydroxybenzoic acid (DHB, 40 mg/ml in 70/30 acetonitrile/water with 0.1% TFA) were applied to the MALDI plate and air-dried. MALDI analysis was performed in reflectron positive mode on an AB SCIEX TOF/TOF 5800 MALDI (Sciex).

### Crosslinking experiments and mass spectrometry analysis of crosslinked peptides

For cross-linking experiments, protein stock was diluted to a final concentration of 50 μM in a total volume of 100 μl using phosphate buffered saline. A 100 mM BS^3^ stock solution was freshly prepared by dissolving 2 mg BS3 in 35 μl Milli-Ǫ water (corresponding to 5 mM final concentration 1:20 stock). For each reaction, 5 μl of BS^3^ stock was mixed with 95 μl of the diluted protein sample (1:100 protein to crosslinker) in an Eppendorf tube and incubated for 1 hour at 4°C. The reaction was quenched by addition of 100 μl of 0.5 M Tris-HCl (pH 7.5) and incubated for 30 minutes at room temperature. For SDS-PAGE analysis, 60 μl of the cross-linked sample was mixed with 20 μl of LDS sample loading buffer (ThermoFisher), heated at 100 °C for 10 minutes, and briefly centrifuged. Subsequently, 30 μl of each sample was loaded onto a gradient gel, electrophoresed at 200 V for approximately 30 minutes, and imaged to visualize cross-linked products. For experiments using DSSO (disuccinimidyl sulfoxide) and DTSSP (3,3′-dithiobis[sulfosuccinimidylpropionate]), reactions were performed under identical conditions. DSSO was used at a 1:100 protein (10 μM):crosslinker (1 mM) molar ratio, while DTSSP was used at a 1:10 protein (5 μM) :crosslinker (50 μM) molar ratio. All reactions were incubated at room temperature for 60 minutes and terminated by the addition of 1 M Tris buffer (pH 8.0) to a final concentration of 20 mM.

The gel bands corresponding to the BCOLg crosslinked using BS^3^, DTSSP and DSSO were removed from the gel, cut into 1 mm³ pieces, and placed into 1.5 ml tubes for destaining, with three incubations (250 μl) in 25 mM NH_4_HCO_3_ containing 50% ACN for 10 minutes at 37 °C. Destained gel bands were dehydrated with 250 μl of pure ACN, then resuspended in 50 μl of 25 mM NH_4_HCO_3_ for proteolytic digestion. The digestion reaction of BCOLg with Trypsin/Lys-C (Promega) at a 1:10 ratio was incubated for 16 hours at 37 °C. Finally, the peptides were desalted using a Pierce C18 spin column (Thermo Scientific) and analyzed by LC-MS (SCIEX TripleTOF 5600) using the SWATH protocol. MS data were analyzed using MeroX 2.0 for crosslinked peptide identification, and the results were visualized in xiView (https://www.rappsilberlab.org/software/xiview/). BCOLg and human genome FASTA-files were used in the crosslinked peptide identification.

### Data availability

All data are available in this manuscript and its supporting information. The structures described in this paper have been deposited with the wwPDB with accession code 9PLV.

## Results

### Purification and enzymatic characterization of recombinant BCOLg for crystallization

Previously, we ^1^ reported the identification of the BCO-like (BCOL) assemblage of CCDs, predominantly in marine invertebrate organisms, suggesting a specialized role in extracellular carotenoid metabolism. To confirm the presence of the BCOLg transcript XP_035666855.1 (LOC118409706 beta,beta-carotene 15,15’-dioxygenase-like) in its expected location, we performed RNAseq analysis of gut tissue of three individual wild-caught lancelets. The LOC118409706 transcript was present in all three specimen guts (247.7±176.2 gene counts) with clear exon expression (Figure 1A) but high density of SNPs. We found 14 amino acid changes with five in asparagines (Figure 1B). The new curated version of BCOLg had eleven amino acid changes (three in the signal peptide) in its sequence compared to the older version of transcript XP_002601288.1. The changes in five of these (181I>L, 395N>S, 466P>A, 454D>A, 473N>S) were confirmed in our transcriptomics data with two in N-linked glycosylation-associated asparagines. We identified five other BCOL transcripts in lancelet gut together with three canonical BCOs transcripts (Table S2). Unlike other metazoan CCDs, BCOL proteins contain signal peptides that facilitate extracellular protein secretion. In lancelet BCOLg, the secretory signal peptide spans amino acid (aa) residues 1 to 22, with a predicted cleavage site between residues 22 and 23 in the sequence: ALS–VT (Figure S1A)^1^.

**Figure 1.**
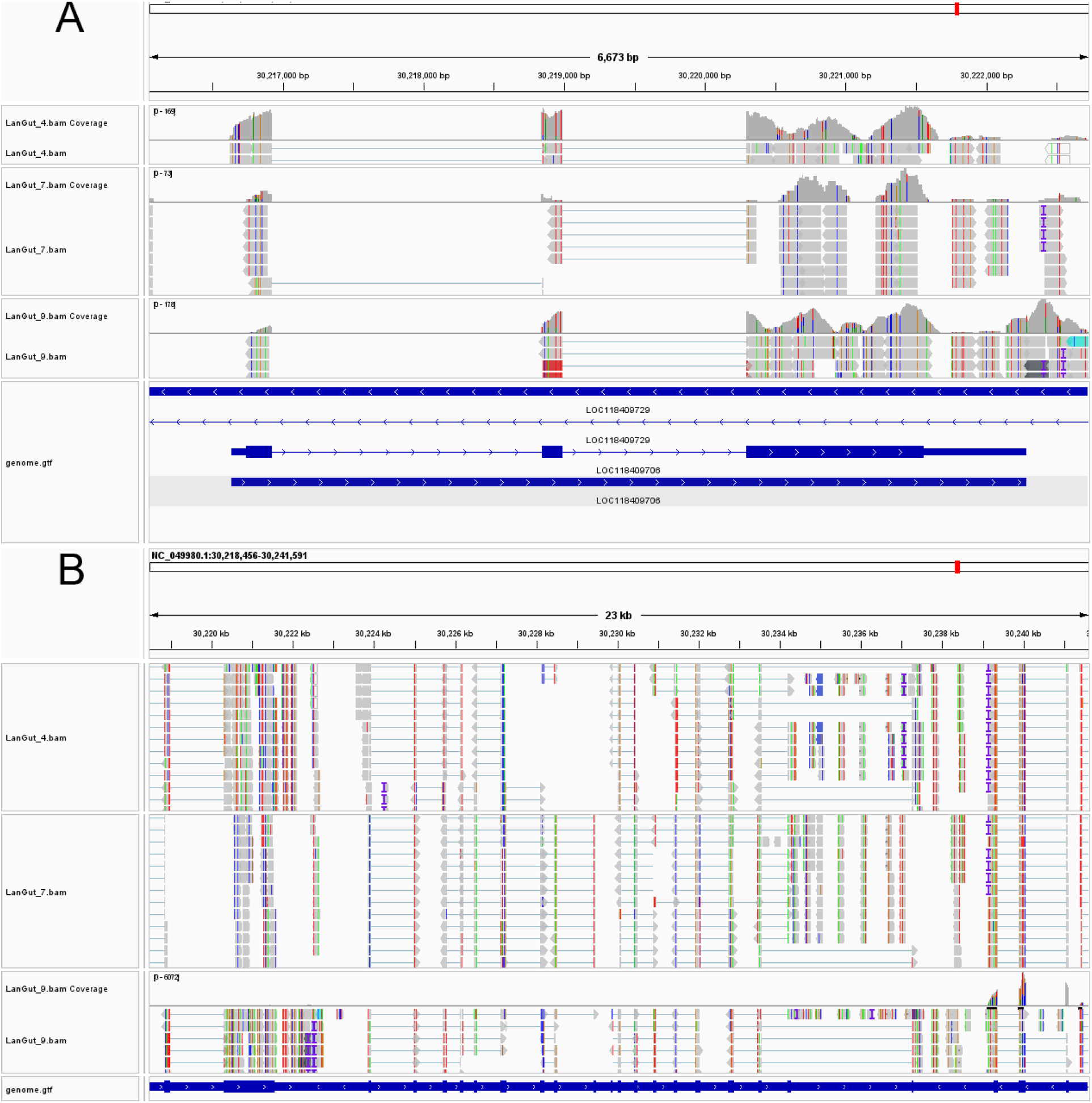
Visualization of BCOLg transcript in lancelet gut RNA-seq data. (A): IGV visualization of full LOC118409706 transcript in all three lancelet samples. The wild caught lancelets were dissected and RNA from gut was isolated. A directional mRNA library (Poly A enriched) was prepared and sequenced with NovaSeq X Plus with depth ∼100M/30G reads per sample (Novogene). (B) IGV visualization of exon 3 in all lancelet samples, with colored SNPs.

In preliminary experiments, BCOLg was secreted into the culture medium when expressed in HEK293F suspension cells^24^. To prepare enough protein for crystallization screening, we expressed BCOLg in Expi293 suspension cells. The culture supernatant was first subjected to PD-10 desalting to remove medium components and to enrich the target protein. The desalted sample was subsequently concentrated then purified by size-exclusion chromatography. Figure S1B illustrates BCOLg purification process. We performed affinity purification of BCOLg using a TALON (cobalt-based) column which led to noticeable protein degradation, likely due to prolonged binding and elution conditions (Figure S2A, B, and C). To improve stability, the purification strategy was shifted to desalting followed by size-exclusion chromatography (SEC), which preserved the integrity and activity of the protein.

The SEC elution profile (Figure 2A) revealed a symmetric peak at approximately 77 ml elution volume with an apparent molecular weight 76kDa, indicating the purified BCOLg exists predominantly as a monomer in solution (Figures 2B and 2C). A small fraction of catalytically inactive (based on *in vitro* assay), aggregated protein eluted in the void volume and was excluded from further analysis. The typical yield was 4-6 mg of purified protein from 120 ml of culture (or 33-50 mg/l culture).

**Figure 2.**
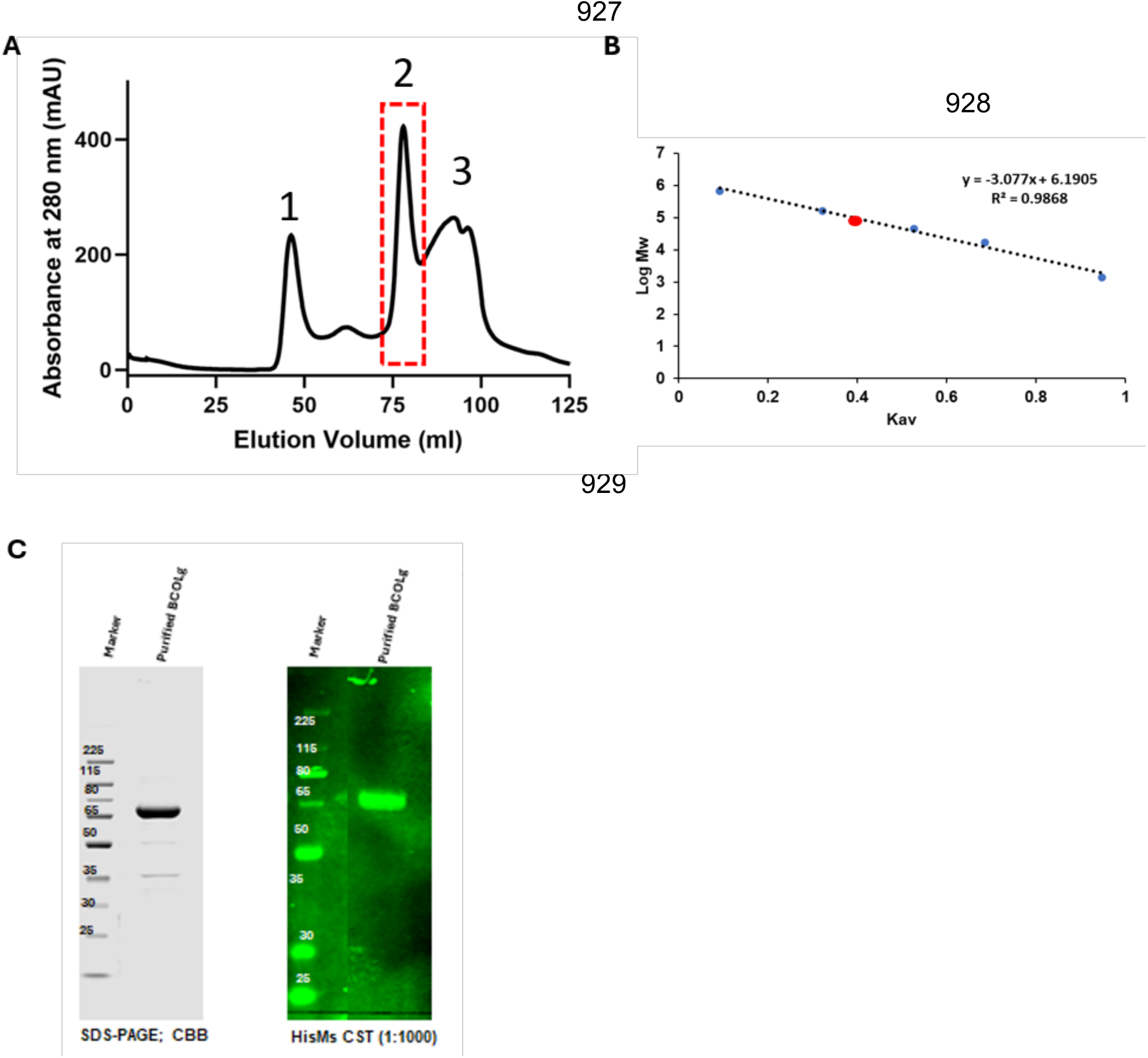
Purification and oligomeric state analysis of recombinant BCOLg protein. (A): Size-exclusion chromatography (SEC) profile showing elution of the purified BCOLg protein, with two major peaks labeled (1) and (2); the main peak (highlighted in red box) was used for further analysis. (B): The calibration curve of log molecular weight (MW) versus Kav indicates that the protein elutes corresponding to an apparent molecular weight of ∼76 kDa. (C): SDS-PAGE analysis (left; Coomassie Brilliant Blue (CBB) staining) shows a predominant BCOLg protein band at the expected size, confirming purity, while western blot (right) using mouse His-tag (HisMs, cell signaling technology (CST) antibody further validates the identity of the purified protein. The partition coefficient (Kav) in gel filtration chromatography describes the fraction of the stationary phase pore volume accessible to a given molecule and is calculated as: 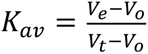 where *V_e_* is the elution volume of the sample, *V_o_* is the void volume determined using a completely excluded molecule, Blue Dextran, and *V_t_* is the total column volume.

No degradation of BCOLg was detected by SDS-PAGE after purification; however, co-elution of the chaperone HSP70 was confirmed by immunoblotting and mass spectrometry (Figure 2C, Figure S3). Fractions corresponding to the entire SEC peak 2 were pooled and analyzed by native PAGE (Figure S2D). Under non-denaturing conditions, BCOLg migrated as a predominant species together with several weaker, slow-migrating bands, suggesting the presence of minor higher-order species or conformationally distinct forms.

### BCOLg crystallization and assessment of its glycosylation

Crystallization trials were performed using the purified catalytically active BCOLg protein under various screening conditions. Although elongated crystals large enough for X-ray diffraction were obtained, they did not yield useable diffraction. As BCOLg is a secreted protein, heterogeneous glycosylation could potentially hinder crystal formation, as well as affect protein stability and solubility. N-linked glycosylation sites are predicted at four residues, Asn98, Asn110, Asn383, and Asn438. To address whether glycans were affecting crystallization, we treated purified BCOLg protein with PNGase F followed by SDS-PAGE. The PNGase F-treated sample exhibited a distinct increase in electrophoretic mobility compared with untreated BCOLg, confirming the presence of N-linked glycans (Figure S4).

To reduce N-glycan heterogeneity and size, BCOLg was expressed in Expi293F GnTI⁻ cells, which lack N-acetylglucosaminyltransferase I (GnTI^−^) activity and express proteins with only high-mannose or hybrid-type N-glycans. BCOLg from Expi293F GnTI⁻ supernatant revealed a reduced glycan complexity compared with protein expressed in parental Expi293F cells, and only simple glycan structure (M5) was identified, as confirmed by mass spectrometry analysis (Figure S4B). MALDI-MS/MS analysis identified only the oligomannosidic N-glycan structure with five mannose residues and two N-acetylglucosamine residues (M5 - Man5GlcNAc2) on BCOLg expressed in Expi293F GnTI⁻ cells as expected in proteins produced in these cells^25^. Subsequent crystallization of BCOLg purified from the Expi293F GnTI⁻ cell line resulted in crystals that diffracted to high resolution, enabling us to solve the protein structure.

### N-Glycosylation affects the enzymatic function of BCOLg in vitro

Next, we investigated the enzymatic activity of purified BCOLg protein. BCOLg (2 mg/ml) purified from Expi293F cells could cleave β-carotene to form vitamin A aldehyde. Activity was ∼32 % of human BCO1 isolated from Expi293F cells (Figure S5A). However, when BCOLg was produced in GnTI^−^ cells with deficient N-glycan production, activity dropped significantly to ∼1.3 % of human BCO1 activity (Figure S5B). This could be explained by decreased solubility of the protein in the assay.

### Crystal Structure of BCOLg

We determined the structure of BCOLg by X-ray crystallography at 2.4 Å resolution. As in other members of the carotenoid cleavage dioxygenase (CCD) family, BCOLg adopts the canonical seven-bladed β-propeller fold, comprising 35 β-strands (Figure 3 and 4). The core of BCOLg consists of seven β-sheets connected by loops, which are interspersed with α-helices that together form a catalytic dome. This dome covers the catalytic iron center and serves as a ceiling over the active site, which is defined by an Fe²⁺ ion held at the propeller axis by four histidine residues (His188, His241, His306, and His500^26^. The bottom face of BCOLg is formed predominantly by the loops connecting the β-sheets and, unlike the dome, contains no significant α-helices.

**Figure 3.**
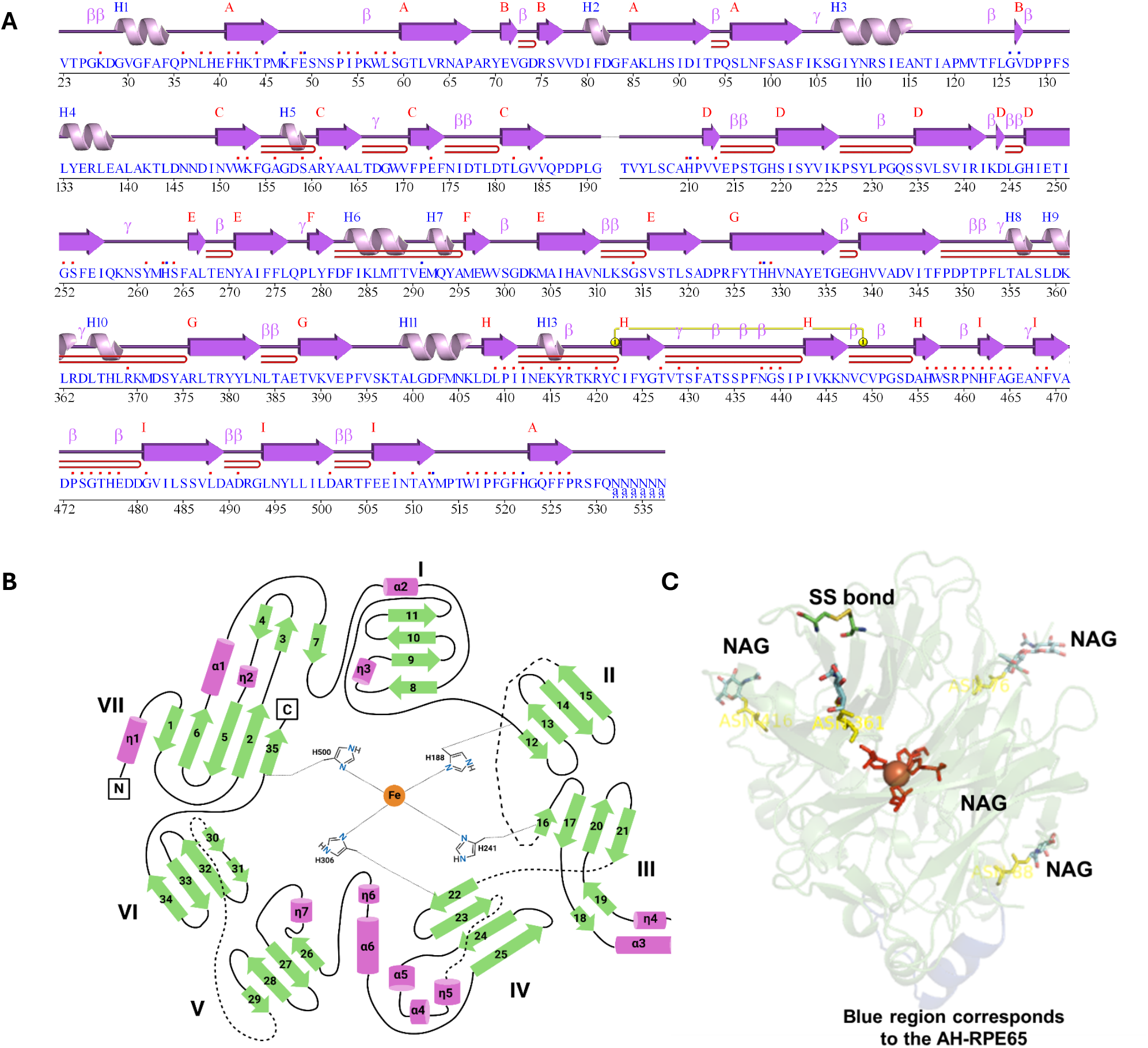
Structural organization and domain architecture of BCOLg protein. (A) The linear schematic depicts the secondary structure elements along the amino acid sequence, including α-helices and β-strands, and connecting loops with key residues and domains annotated. Yellow line indicates a disulfide bond stabilizing the protein structure. Because BCOLg is a secreted protein whose first 22 amino acids constitute the signal peptide (see Figure S1), residue numbering begins at position 23 of the mature protein, in accordance with the convention of numbering based on the gene sequence rather than the mature protein. (B) Topological diagram illustrating the arrangement of β-sheets and α-helices organized into distinct structural domains (I-VII), with key residues and the central metal-binding histidine residues highlighted. (C) Three-dimensional structural model displaying overall fold, N-linked glycosylation sites (NAG), and disulfide (SS) bond; highlighted regions indicate important structural features within the protein. In both (B) and (C) the Fe²⁺ ion is depicted as an orange sphere.

**Figure 4.**
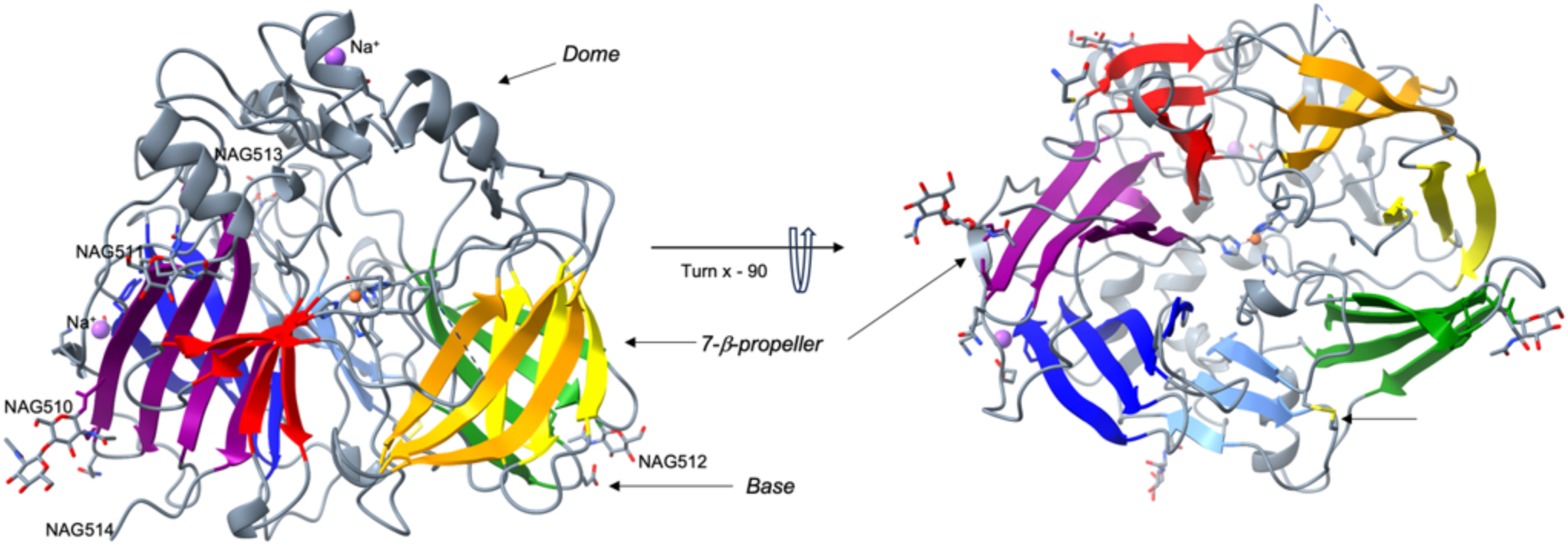
Two views of the seven-bladed β-propeller structure of BCOLg, with blades colored as follows: Blade I, red (β7–β12); Blade II, orange (β13–β16); Blade III, yellow (β17– β21); Blade IV, green (β22–β26); Blade V, blue (β27–β31); Blade VI, indigo (β32–β34); and Blade VII, violet (β1–β6, β35). The loops connecting the blades and the catalytic cap are shown in gray, along with the glycans at Asn98 (NAG510, NAG514), Asn110 (NAG511), Asn383 (NAG512), and Asn438 (NAG513), and Cys422 and Cys449 disulfide bond shown in yellow. The Fe²⁺ ion, coordinated by four histidine side chains, is depicted as an orange sphere, and the Na⁺ ions from the crystallization solution are colored magenta.

The asymmetric unit contains two BCOLg molecules arranged base-to-base. The A/B interface (Figure S6A) comprises seven hydrogen bonds and buries a total surface area of 1200 Å² (600 Å² + 610 Å²). The second large interface B/A in the crystal is formed between molecule A and a symmetry-related molecule B distinct from the one in the asymmetric unit (Figure S3B); it corresponds to a side-to-side packing of two seven-bladed β-propellers contacting each other through blades VI (indigo) and VII (violet), comprises six hydrogen bonds, and buries a total surface area of 1480 Å² (740 Å² + 740 Å²). In both cases, the hydrogen bonds reflect close crystal packing, in which protruding side chains interact with their immediate environment, rather than specific, function-related contacts. The base-to-base arrangement of the two BCOLg monomers in the asymmetric unit differs from those observed in structurally related proteins, such as ACO (PDB ID 2BIW), in which the hydrophobic patches face each other (buried surface area (BSA) 1370 Å²), and RPE65 (PDB ID 3FSN), in which the entrances to the active sites face the same way (BSA 3100 Å²).

The two most suitable reference structures, due to their homology and structural likeness to BCOLg, are the retinal-forming ACO and membrane-associated RPE65 retinol isomerase. BCOLg superimposes ACO and RPE65 with a root mean square deviation (RMSD) of 3.8 and 3.5 Å for 463 and 449 matched Cα positions, respectively. 2BIW RMSD between 275 pruned atom pairs is 1.09 Å; (across all 463 pairs: 3.8) 3FSN: RMSD between 307 pruned atom pairs is 1.05 Å; (across all 449 pairs: 3.5). Four N-acetylglucosamines (NAGs), at Asn98, Asn110, Asn383, and Asn438, were resolved (Figures 3), not seen previously in CCDs. As a native secreted protein, BCOLg possesses a disulfide bond between Cys422 and Cys449 (Figure 3 and 4). This feature has not been reported for intracellular BCOs; even when intracellular BCOs have two nearby Cys residues, their geometry does not favor covalent bond formation.

### Structural motifs and active site

BCOLg shares most features with ACO except for the extensive loop between β24 and β25 of blade IV present in RPE65 (L372-T392), *C. elegans* rBCMO-1, and rBCMO-2 (PDB IDs 7WH0 and 7WH1) (Figure 5A) ^8^. BCOLg and ACO have a blade IV loop shorter than the 29 aa loop 371-400 of RPE65 (Figure S7, Figure 5A). The BCOLg electron density revealed that loops 191-203 and 232-233 were disordered but were modeled by AlphaFold (Figure 5B and C). While the residues 232-233 are modeled with reasonable confidence (Figure 5B), the position of residues 191-203 is predicted with only moderate reliability due to the high mobility of the loop (Figure 5C). Furthermore, BCOLg does not have the conserved -PDPCK-motif, which instead is replaced by a -VDPPF-motif. While the -PDPCK-motif is not resolved in native RPE65, it was successfully resolved in the putative nematode rBCMO-1 and rBCMO-2 ^8^. Superposition of BCOLg and rBCMO-1 shows that the -PDPCK-motif, which plays a key role in post-translational palmitoylation of vertebrate BCOs, is substantially shifted relative to the corresponding loops in BCOLg and ACO (Figure 5D and 5E). Consistent with it being a secreted protein, BCOLg displays a disulfide bond between Cys422 and 449, absent in cytosolic RPE65 and ACO (Figure 5F).

**Figure 5.**
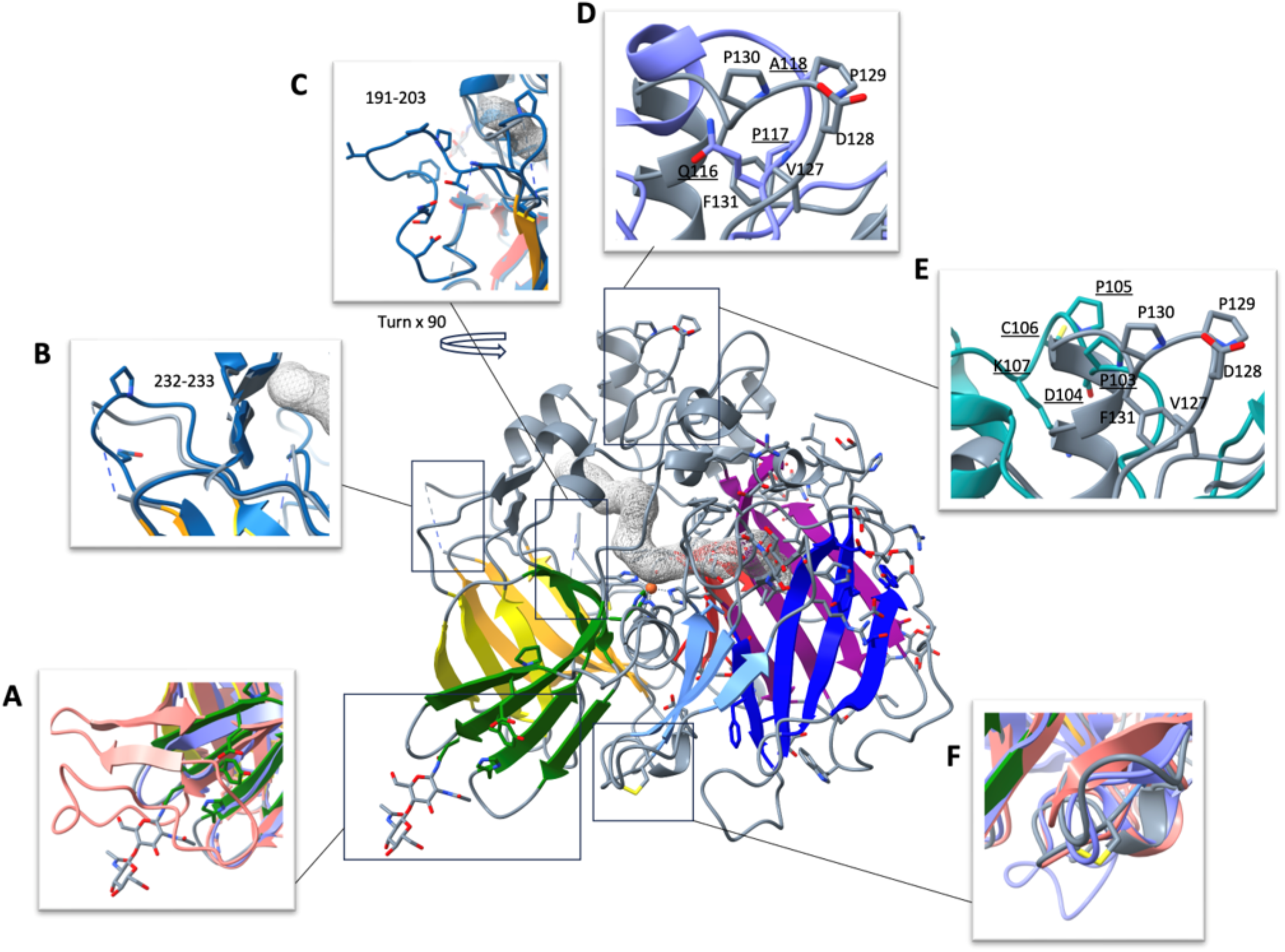
Summary of BCOLg structural features compared with the AlphaFold model (blue), BCMO1 (teal), ACO (2BIW, violet), and RPE65 (3FSN, salmon). (A) BCOLg and ACO (violet) have a shorter blade IV loop (residues 383-388 of BCOLg) than the 29-residue loop (residues 371-400) of RPE65 (salmon). (B and C) BCOLg structure superimposed on the AlphaFold model (blue), showing the positions of the missing residues. Residues 191–203 are predicted with only moderate reliability, owing to the high mobility of this loop. (D and E) In BCOLg (gray) and ACO (violet), the -PDPCK-motif is replaced by the sequences -VDPPF- and -ǪPAGG-, respectively, whereas in the C. elegans putative BCMO1 (teal), and bovine RPE65 the motif is shifted. Residues are labeled in plain type for BCOLg and underlined for ACO and C. elegans BCMO1. (F) A disulfide bond between Cys422 and Cys449 is present in BCOLg (gray) but absent in ACO (violet) and RPE65 (salmon).

A hydrophobic tunnel leads from the protein exterior to the iron-defined active site, enabling carotenoids to pass to the catalytic center, consistent with other BCOs (Figure 5A). Adjacent to this tunnel, BCOLg has a nonpolar surface patch consisting primarily of protruding leucines and tyrosines. It was proposed that ACO uses this patch to dip into the membrane to extract its nonpolar substrate from there. The nonpolar patch of BCOLg is much smaller than ACO’s and consists of Leu133, Tyr134, Leu137, Leu140, Tyr229, Leu230, and Ile284.

The propeller’s top face, defined by the segments connecting the outer strand of one sheet with the inner strand of the next, is covered by a helical cap housing the active site. The Fe^2+^ ligand is located near this top face on the propeller axis. It is coordinated by four His residues. In the case of both ACO and RPE65 the positions of the three of these His are stabilized by direct or water-mediated hydrogen bonds with three Glu residues (Glu150/148, Glu370/417, and Glu426/469 in ACO/RPE65), forming a second coordination shell around the iron. BCOLg is unique among CCDs in that two of these glutamates are replaced by Asp167 and Asp408, which occupy the same positions and similarly stabilize the iron-coordinating histidines (Figure 6). A direct hydrogen bond forms between His263 and Asp167, enabled by a ∼2 Å shift of the main chain of the latter towards the histidine residue. In contrast, ACO’s His238 is shifted closer to Glu150 for a direct hydrogen bond between them, while RPE65’s Glu148 and His241 form a water mediated hydrogen bond stabilized by Tyr239^9,27^. The position of Asp408 closely coincides with Glu386 (ACO) and Glu417 (RPE65), and all three form a direct hydrogen bond with His328 and its respective analogs (Figure 6B). The stretch 405-422 containing Asp408 is anchored to the base of β27 strand by the somewhat distant disulfide bond, and Asp408 main chain position is further stabilized by hydrogen bonding with Thr427 of β27.

**Figure 6.**
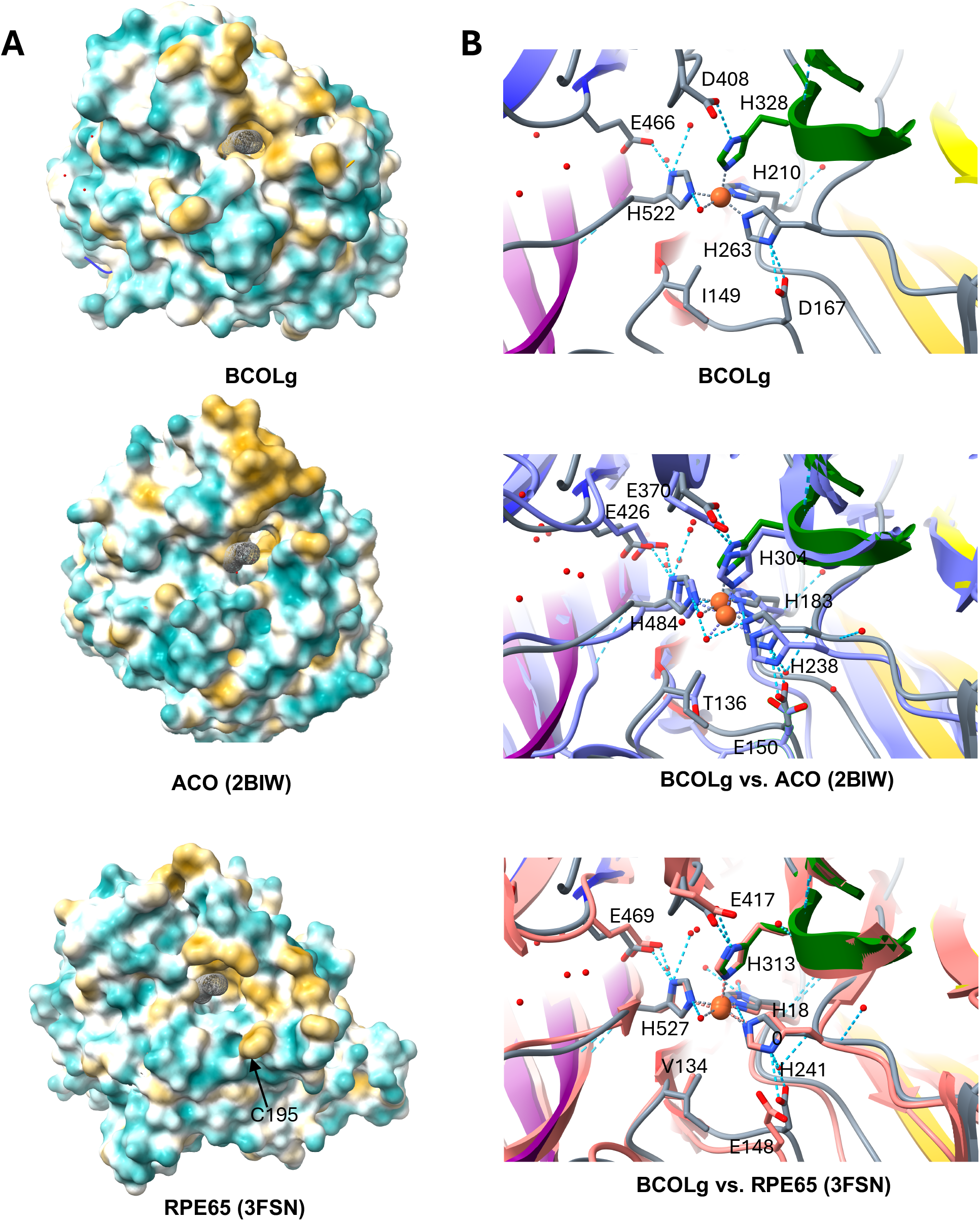
The hydrophobic patch at the catalytic tunnel entrance and the active site arrangement in BCOLg, ACO, and RPE65. (A) Hydrophobicity of the residues lining the tunnel entrance (black mesh), with hydrophobic residues shown in yellow and hydrophilic residues in cyan. (B) Active sites of BCOLg, ACO (2BIW, violet), and RPE65 (3FSN, salmon), showing the three Fe²⁺-coordinating histidines of the first coordination sphere and the three glutamate/aspartate residues that stabilize them through direct or water-mediated hydrogen bonds, forming the second coordination shell around the iron. The Fe²⁺ ion is depicted as an orange sphere.

### Analysis of Substrate-Accommodating Channel

Using the MOLEonline tool ^28^, we modeled the substrate-accommodating channels of BCOLg, ACO, and RPE65 (Figure 7A). The inner surface of the BCOLg channel is defined by the following residues: Phe34, Phe81, Phe124, Glu166, Tyr261, Thr327, Leu409, and Thr429. In the case of ACO, it is formed by Tyr24, Phe69, Phe113, Trp149, Phe236, Phe303, Phe371, and Leu400, whereas for RPE65 it is lined by Phe16, His59, Phe61, Gly104, Thr147, Tyr239, Phe312, Phe418, and Gly438. Superposition of corresponding residues (Figure 6B) shows that the overall channel geometry of BCOLg is less restricted than in ACO and RPE65. Four positions are hypothesized to be critical for the channel’s geometry and, thus, its specificity: Phe81, Phe124, Thr166, and Leu409. A less restricted channel suggests BCOLg may have a lower catalytic efficiency or broader substrate specificity or may be able to accommodate bulkier substrates compared to ACO and RPE65, whose tighter channels might impose greater selectivity.

**Figure 7.**
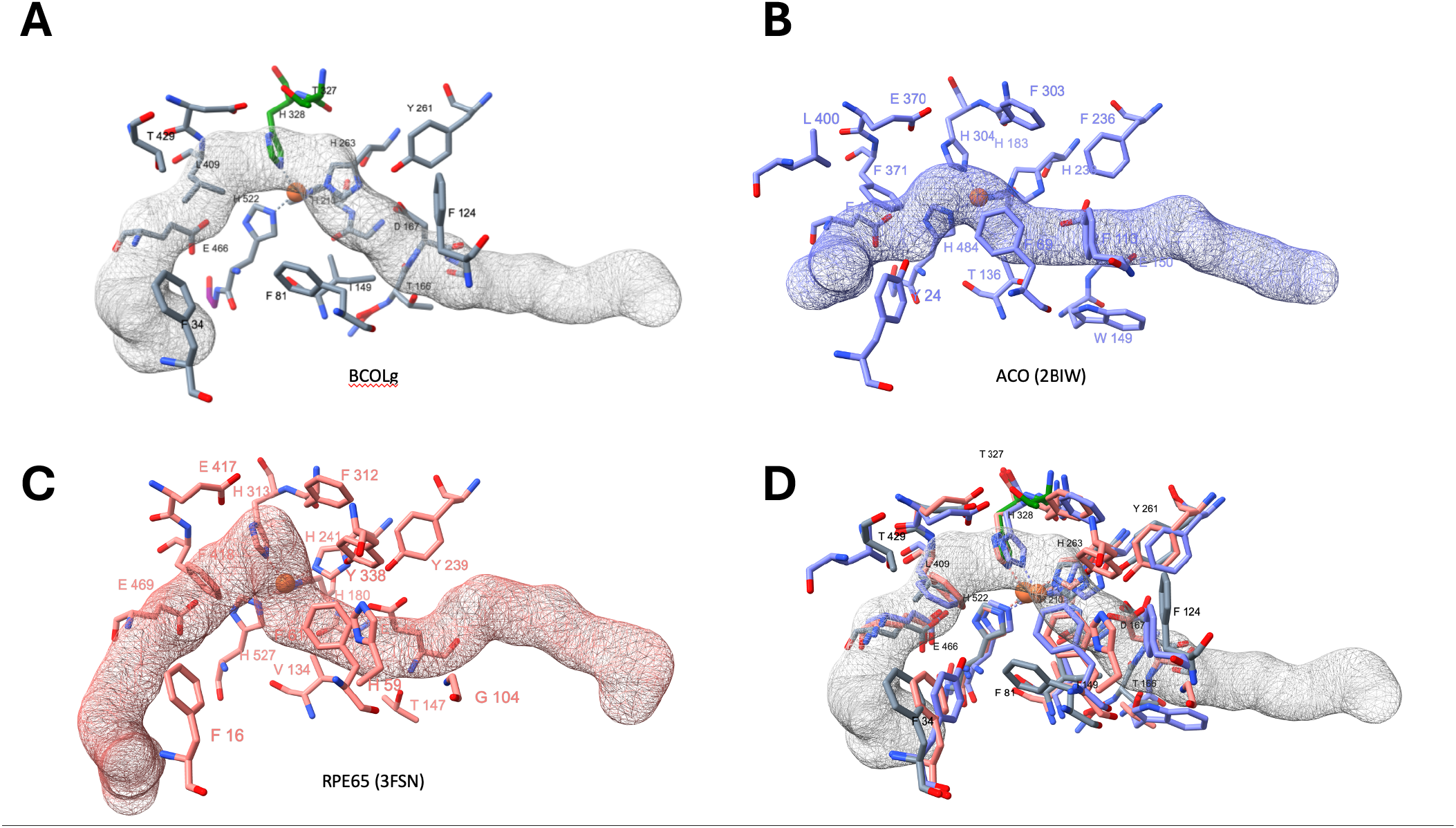
Catalytic tunnels of BCOLg, ACO, and RPE65. (A–C) Individual catalytic tunnels modeled with the MOLEonline tool. Hydrophobic residues lining the substrate-binding channel are shown explicitly, and the mesh depicts the overall shape and size of the channel. (D) Superposition of the channel-lining residues of BCOLg, ACO, and RPE65 onto the BCOLg channel reveals a less restricted geometry than in ACO and RPE65, in which several bulky side chains would clash with a substrate occupying the channel. The Fe²⁺ ion is depicted as an orange sphere.

### Additional electron density

BCOLg crystals demonstrated the presence of a weak, tube-shaped electron density feature within the active site. This density was observed with added substrate and even in the absence of a deliberately added substrate, and its appearance is highly reminiscent of similar features reported for its structural homologs, including ACO (PDB IDs 2BIW, 4OU8, 4OU9) and RPE65 (PDB 3FSN). Experiments to visualize a substrate in the structure did not reveal electron density in the binding tunnel. Further experiments are in progress. Solvent density was observed in the binding tunnel of the structure. The origin of similar density in earlier CCD studies was hypothesized to arise from a component of the crystallization mixture ^29^. Due to the weak signal strength and the poorly defined shape of the feature in the BCOLg map, it was left unattributed in the final model, consistent with best practices for crystallographic data interpretation (data not shown).

### Far-UV CD Reveals a Conserved Native Fold of BCOLg

To examine the effect of buffer composition and expression system on the secondary structure, far-UV CD spectra were recorded for Expi293-derived BCOLg in Tris buffer containing glycerol (Figure S8, Protein 1), the same Expi293-derived BCOLg after dilution into Tris buffer without glycerol (Figure S8, Protein 2), and GnTI^−^-derived BCOLg in Tris buffer without glycerol (Figure S8, Protein 3). Although both Expi293- and GnTI^−^-derived proteins were purified by gel filtration in Tris buffer containing glycerol, glycerol was removed from selected samples prior to CD measurements. The overall spectral profiles were consistent with a predominantly β-sheet secondary structure, as expected for the seven-bladed β-propeller scaffold of BCOLg. While all three samples retained the characteristic features of a folded β-sheet-rich protein, differences in spectral intensity were observed (Figure S8, top panel). Removal of glycerol from the Expi293-derived BCOLg resulted in a modest increase in the negative ellipticity, whereas the GnTI^−^-derived protein exhibited the largest negative ellipticity among the three preparations. Projection of the [Δε]_222nm_ vs [Δε]_200nm_ onto the Protein Circular Dichroism Data Bank (PCDDB) reference map ^30,31^ (Figure S8, bottom panel) placed all samples within the pre-molten globule to native folded region, indicating that the overall structural integrity of BCOLg was maintained under all conditions. These results suggest that glycerol and the expression system influence the conformational ensemble or structural dynamics of BCOLg without altering its overall native fold.

### Oligomeric State of BCOLg Protein

The crystal structure of BCOLg revealed two monomers packed in the asymmetric unit (Figure 3), suggesting that the protein may form a dimeric assembly under crystallization conditions. To determine whether this dimerization is physiologically relevant or an artifact of crystal packing, we examined the oligomeric state of BCOLg using a combination of biophysical and biochemical methods.

First, size-exclusion chromatography (SEC) was performed to assess the apparent molecular mass of BCOLg in solution. The elution profile showed that the protein migrated primarily as a species with a molecular weight around ∼76 kDa whereas the calculated molecular weight of protein is approximately 63-64 kDa. Consistent with its glycoprotein nature, the slightly higher apparent molecular weight observed by SEC is consistent with the increased hydrodynamic radius of the glycosylated (Figure 2B). A minor fraction eluted near the void volume, representing a small population of higher-order aggregates, as expected.

To further characterize this behavior, we analyzed BCOLg under non-denaturing conditions using native PAGE. Native PAGE analysis revealed slowly migrating species, suggesting either transient oligomerization or conformation heterogeneity that alters electrophoretic mobility rather than stable high-order oligomers.(Figures S2D).

To explore this observation, we performed chemical crosslinking experiments using three amine-reactive crosslinkers of varying spacer arm lengths and chemical properties— bis(sulfosuccinimidyl)suberate (BS3), 3,3′-dithiobis (sulfosuccinimidyl propionate) (DTSSP), and disuccinimidyl sulfoxide (DSSO). Upon crosslinker treatment, a shift in the molecular weight from monomeric (∼60 kDa) to dimeric (∼120 kDa) BCOLg was observed by SDS-PAGE (Figure 8A). The persistence of a monomeric population likely reflects incomplete crosslinking, possibly due to limited crosslinker concentration or steric hindrance at the reactive lysine residues. These findings further supported the presence of dimeric interactions under certain conditions.

**Figure 8.**
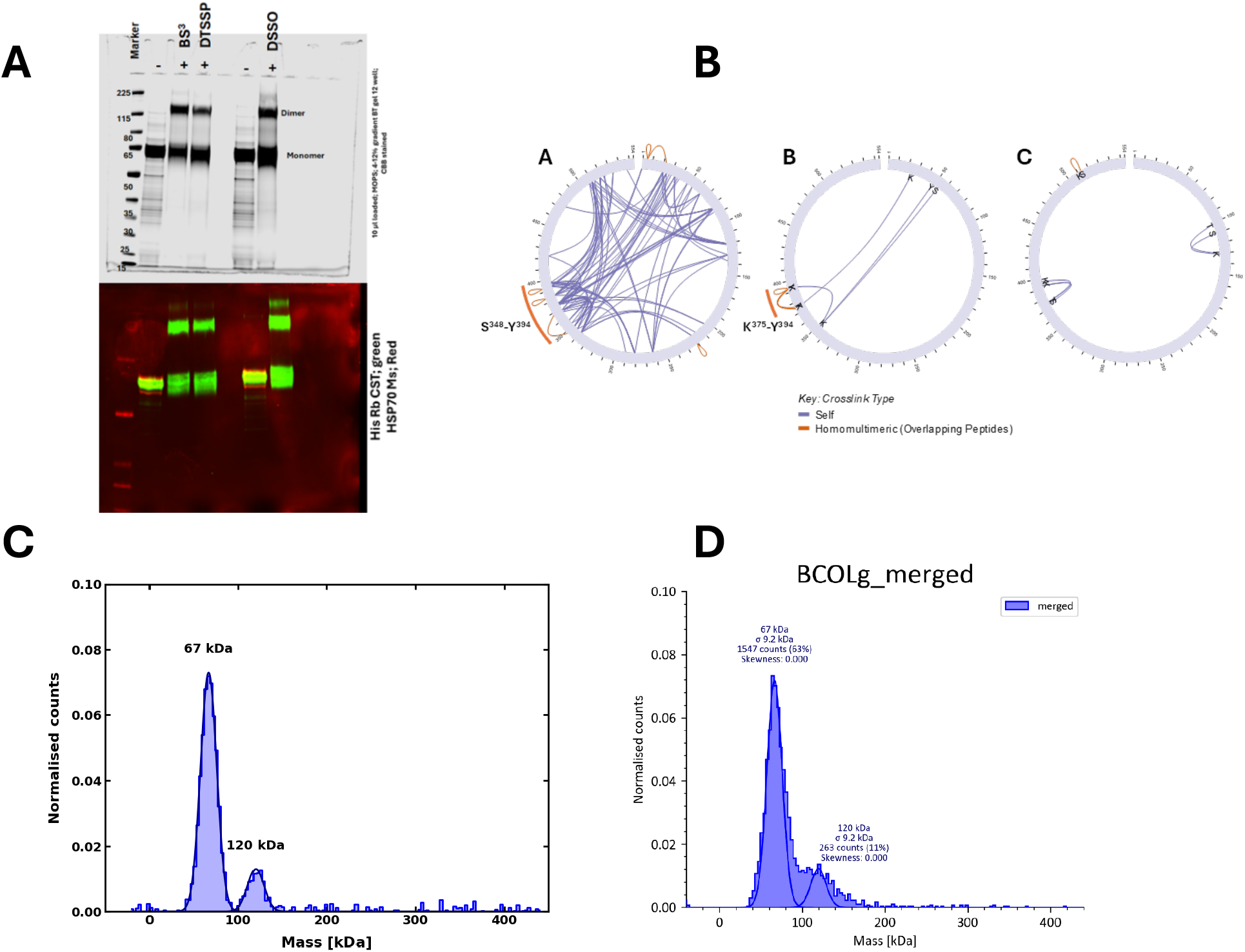
Biochemical and crosslinking analyses reveal oligomerization and interaction architecture. (A) SDS-PAGE (top) and western blot (bottom) of BS³-, DTSSP- and DSSOtreated BCOLg samples show crosslinked species corresponding to monomer and dimer forms. (B) Circular crosslink maps depicting intra- and inter-molecular crosslinks respectively. (1) Extensive self and overlapping (homomultimeric) crosslinks indicate a dense interaction network. (2) Reduced crosslinking highlights specific residue–residue contacts. (3) Minimal crosslinks suggest limited or transient interactions. Crosslink types are indicated as self-links (gray) and homomultimeric/overlapping peptides (orange). (C-D. C. fitted mass, D. raw mass photometry data). Mass photometry analysis of diluted (0.1 mg/mL) solution of purified BCOLg demonstrated mass distribution of detected peaks at ∼67 kDa (63%) and ∼120 kDa (11%), consistent with predominant monomeric and low abundance dimeric forms, respectively.

When crosslinked peptides were analyzed by LC-MS several homomultimeric crosslinked peptides (two different BCOLg molecules) between K375-Y394 and Y394-K393 in BS3 (11.4 Å) crosslinking, and S348-Y394 K393/Y394-K393/Y394, and K384-K384 in DSSO (10.3 Å) crosslinking, were revealed. No crosslinked peptides were detected in this region using the longer arm DTSSP (12 Å), indicating that the distance between the two BCOLg molecules in this area ranges from 10.3 to 11.4 Å, based on the spacer arm lengths for BS3 and DSSO. Additional inter-BCOLg crosslinked peptides were identified between K5-K21/K22 K221-K221, and K221/T228)-K221/T228in DSSO crosslinking and between S507 K510 in DTSSP crosslinking. In addition, intramolecularly crosslinked peptides were identified for all crosslinking reactions, mainly with DSSO, as this molecule is suitable for intramolecular crosslink analysis (Figure 8B).

Given the indications of concentration-dependent dimerization, we next employed mass photometry to assess the oligomeric state of BCOLg at nanomolar concentrations (Figure 8C). This single-molecule technique revealed that BCOLg is predominantly monomeric in dilute solution, with only a small fraction corresponding to the dimeric species. The relative abundance of the dimer increased slightly with protein concentration, confirming that the association is weak and reversible.

Taken together, these results indicate that BCOLg exists primarily as a monomer in solution but can form dimers under concentrated conditions, as seen in the crystal lattice, native PAGE, and crosslinking assays. The observed dimerization is therefore likely concentration-induced, rather than reflecting a physiologically stable state. These findings suggest that BCOLg functions as a monomer under physiological conditions, with dimer formation being a transient or crystallization-induced phenomenon. DLS of BCOLg protein used for crystallization had hydrodynamic diameter of 6.2±2.4 nm and PDI of 0.26 by DLS (Figure S9) corresponding to theoretical hydrodynamic diameter of a monomer (5.93 nm)^32^ and close to the 5.7 nm diameter obtained from the X-ray crystal structure reported here (PDB ID: 9PLV).

### Structural analysis of BCOLg adjacent monomers interface in crystal structure

To understand the structural basis of the possible functional oligomerization, as was previously shown for other CCDs (BCMO1 and RPE65)^10,29,33^ we analyzed the adjacent monomer’s interface by PDBSUM analysis. In the BCOLg crystal structure (Figure S10A), chains A and B formed an interface encompassing 14 and 15 residues, with a BSA of ∼666 Å² per monomer. The interface is stabilized by four hydrogen bonds and 71 non-bonded contacts. The predominant interactions involved Ser529 and Gln531 (chain A) engaging with Asn270 and Glu334 (chain B), forming a conserved Ser-Glu and Gln-Asn/Glu polar interaction network. These contacts contribute to a moderately extensive and flexible interface stabilized primarily through hydrogen bonding and hydrophobic packing. Additionally, we crystallized BCOLg in the presence of its quasi-substrate, 8-apocarotenal. However, due to the limited solubility of the latter, we obtained a low occupancy substrate analog-bound form with nearly identical crystal structure with lower interface area of ∼BSA 540 Å² per monomer, but we were unable to reliably see any quasi-substrate molecule (data not shown).

## Discussion

Here we present the structure of a novel carotenoid cleavage dioxygenase (CCD) of lancelet *(Branchiostoma fforidae;* BCOLg) at 2.4 Å resolution, with several features not heretofore seen in previously crystallized CCDs. The presence of BCOLg transcript in the gut of animals was confirmed by RNAseq gene counts analysis and we propose that the BCOLg protein is secreted in lancelet gut for carotenoid cleavage and/or vitamin A synthesis there. CCDs are a diverse family of enzymes of crucial importance, not least of all in human physiology and health. The BCOLg structure maintains the canonical 7-bladed propeller structure of CCDs and stilbene-cleaving oxygenases with a non-heme ferrous ion coordinated by 4 histidines. The novel features include a secretory signal peptide, the presence of N-linked glycosylation (four in BCOLg), an intramolecular disulfide bond, and the replacement of two of three second-shell fixing Glu residues with Asp residues (substitution of E to D appears to be general feature of BCOL proteins (Table S2), while lacking the “metazoan” loop and the -PDPCK-motif common to all other metazoan CCDs. These features expand the repertoire not only of metazoan CCDs but of CCDs/SCOs across all kingdoms of life. We propose that BCOLg is a member of a basal chordate expansion of an ancestral sister clade of metazoan CCDs, and as such, an important aspect of CCD evolution. This may be analogous to the later expansion/specialization of metazoan CCD paralogs, especially of RPE65s, within the vertebrate lineage.

Artificial secretory signal peptides are added to many sequences to generate proteins secreted into media to facilitate purification for crystallization ^34,35^. When we first identified them, we noted that BCOL proteins possessed secretory signal peptides^1^ and, accordingly, we expressed BCOLg as a secreted protein. With the knowledge that this signal peptide can successfully export a 7-bladed propeller fold protein out of eukaryotic cells, our ongoing goal is to employ this naturally occurring secretory signal sequence of BCOLg for expression of other animal CCDs, including pathological and other mutants of RPE65. Relatively few metazoan CCDs and no vertebrate recombinant CCDs have been crystallized. RPE65 was originally crystallized from an abundant native source^9^; no such source exists for BCO1 or BCO2. When recombinant CCDs are overexpressed, prokaryotically or eukaryotically, they are prone to significantly co-express with the relevant chaperone proteins. This is a serious practical problem in purification as the general size of chaperones is the same as that of most CCDs. We hypothesize that expression of CCDs using this secretory signal peptide via secretion from mammalian cells will be a practical solution to this problem.

An important structural feature not present in BCOLg is the aa371-400 loop (in RPE65; the “metazoan loop”) of animal CCDs. BCOLg and ACO have a blade IV loop shorter than RPE65’s 29-residue aa371-400 loop. BCOLg also lacks the -PDPCK-motif present in other metazoan CCDs. This is contained within a ∼30 residue intrinsically disordered region that is not resolved in RPE65 crystal structures, part of which can fold into an amphipathic α-helix in the presence of membranes^36,37^. The homologous -VDPPF-motif found in BCOLg does not contain Cys, has much weaker amphipathic α-helix properties and presents as a well-resolved α-helix in our BCOLg structure.

The presence of four N-linked glycosylation sites on BCOLg, at Asn98, Asn110, Asn383, and Asn438, is another novel feature of the BCOLs among CCDs. While not unexpected for secreted extracellular proteins, it was initially problematic as crystals of the fully glycosylated BCOLg did not diffract adequately. We circumvented this by expressing BCOLg in Expi293F GnTI^−^ cells deficient in N-acetylglucosaminyltransferase I^38^. This limits N-linked glycosylation to homogeneous Man_5_-GlcNac_2_, amenable to structural studies. Interestingly, b-carotene cleavage activity of the GnTI^−^ BCOLg was significantly reduced compared to the fully glycosylated form. Transcriptomic analysis demonstrated high variability in asparagine residues (changing to serine) in wild-caught animals (4-6 possible asparagine N-glycosylation sites). This may be explained by the fact that *Branchiostoma* is known to have a exceptionally high genomic heterozygosity that appears to be driven by a high rate of germline de novo mutation.^39^

The crystal presents BCOLg as a dimer packed in an asymmetric unit, questioning the physiological relevance of the dimer. We studied the oligomeric state of BCOLg to address this uncertainty. Size exclusion chromatography, cross-linking, and mass photometry experiments indicated that BCOLg is a monomer in solution but subject to a concentration dependent dimerization similar to detergent-induced oligomerization for BCO1^33^. Furthermore, the presence of an intramolecular disulfide bond in BCOLg, between Cys422 on the loop just prior to β−strand #27 and Cys449 on the loop just after β−strand #28, a feature not seen in other CCDs, may be important in dimerization. Why it is required for three-dimensional structural stability in BCOLs, but not in intracellular CCDs is not known, especially as the location of the disulfide bridge is in a place not necessarily requiring extra stability. It is, however, on the dimerization face of the monomers adjacent to the polyanionic signature -EDDGV/I/Y-motif common to all CCDs. Perhaps local stabilization is necessary in this region to facilitate potential dimerization.

The replacement of two of the fixing second-shell Glu residues, that are otherwise canonical in CCDs, by Asp residues is another remarkable feature of BCOLs. In protein structures in general, disparate second-shell residues, distal residues not in contact with substrate, are important for stabilizing the first-shell residues that are crucial to catalytic sites, coordination of metal ligands, etc. Substituting second-shell residues in various enzymes can have marked effects on substrate affinity, catalytic efficiency^40^. Electrostatic perturbations, such as altered pKa values or protonation equilibria of catalytic residues, can result from substitution of important second-shell residues^41,42^. In the oxygen-evolving complex (OEC) of the Photosystem II of plants, algae, and cyanobacteria, for example, second-shell residues play an essential role in transporting protons and oxygen out of the OEC while allowing access to water^43^. Even in transport proteins like transferrin, second-shell residues (e.g., the “dilysine trigger”) play an important role in iron binding/release^44^. It is clear that Glu and Asp residues in a sequence are not always directly interchangeable, and this is the case in CCDs. We and others have found that replacing paralogous second-shell Glu residues in vertebrate RPE65 (E148 and E417 in RPE65, and E405 in mouse BCO1) severely affects catalytic activity^45–48^. Indeed, the missense variants E148D and E417Ǫ in human RPE65 are known pathogenic mutations^45,48^. In particular, Nikolaeva et al.^46^ suggested that when Asp replaces Glu417 in RPE65 it is too distant (∼5.7 Å) to form a hydrogen bond with H313, potentially precluding an important electrostatic interaction required for coordination of Fe^2+^. In BCOLg, there is 2.8 Å between D408 and H328 (paralogous to the E417-H313 pair of RPE65) and a hydrogen bond is formed. In general, Asp residues in proteins have a lower pK_a_ (4.1 in acidic proteins and 3.3 in basic proteins; lowered with increased number of intramolecular hydrogen bonds) than Glu (∼4.3 in acidic or basic) that are not markedly affected by the number of these bonds^49^. This would indicate a specific effect of the replacement of the two second-shell Glu residues by Asp residues in BCOLg.

We hypothesize that BCOL proteins are secreted into the lancelet gut and probably mixed with its algal diet along with other digestive enzymes. We show the presence of six (out of seven annotated) BCOL transcripts including BCOLg in the gut of the lancelet by RNAseq. We speculate that algal carotenoids are oxidatively cleaved by the BCOL enzymes and potentially used as a carbon source for the organism’s metabolism. During the digestive process food is solubilized in the intestine, explaining why these proteins do not need to reach into membranes to acquire hydrophobic substrates like many other CCDs. It is known that in other aquatic organisms carotenoids are broken down during their digestive processes^50^. When compared to ACO^29^ and RPE65^51^, BCOLg’s less restricted channel suggests that BCOLg may have a lower catalytic efficiency or broader substrate specificity or may be able to accommodate bulkier substrates compared to ACO and RPE65, whose tighter channels reflect greater selectivity for particular substrates. Given BCOLg’s speculated role as an enzyme involved in the digestive process of lancelet, we favor the possibility of a broader substrate specificity. Additionally, we also found that the nonpolar surface patch, consisting mostly of protruding Leu and Tyr residues, adjacent to the substrate tunnel is much smaller than the equivalent patch in ACO. As we postulate, BCOLg does not need to dip into membranes to acquire its substrate, as ACO does, this may explain the smaller size of the patch.

In conclusion, we have established structural parameters of a new subfamily of metazoan CCDs. While many of the features identified in BCOLg are of particular relevance to its organismal and postulated physiological roles, some of them can be leveraged to investigate CCDs more difficult to purify that are directly implicated in human physiology and health. With this successful crystallization, we plan to use BCOLg as a platform to model structural aspects of RPE65 and other vertebrate CCDs in an easily accessible secreted protein format. Such studies are in progress.

## Acknowledgements

The authors wish to acknowledge the assistance and guidance of Dr. Susan Gentleman and Dr. Vijayasarathy Camasamudram in the original conception of this study. We thank Grzegorz Piszczek and Di Wu (Biophysics Core Facility, NHLBI) for assistance with the mass photometry experiments. This work was supported by the Intramural Research Program of the National Institutes of Health (NIH). The contributions of the NIH authors were made as part of their official duties as NIH federal employees, are in compliance with agency policy requirements, and are considered Works of the United States Government. However, the findings and conclusions presented in this paper are those of the authors and do not necessarily reflect the views of the NIH or the U.S. Department of Health and Human Services.

## Supporting data

**Table S1.** Data collection and refinement statistics. (*Highest resolution shell is shown in parenthesis.)

| **PDB ID** | **9PLV** |
|---|---|
| **Data collection** |  |
| Space group | C*2* |
| Cell dimensions |  |
| *a*, *b*, *c* (Å) | 98.46, 139.91, 93.76 |
| α, β, γ (°) | 90, 98.26, 90 |
| Resolution (Å) | 24.85 - 2.40 (2.46 - 2.40) * |
| *R*<sub>sym</sub> or *R*<sub>merge</sub> | 0.089 (0.328) |
| *I* / σ*I* | 11.98 (2.82) |
| Completeness (%) | 98.1 (89.6) |
| Redundancy | 4.25 (3.0) |
| **Refinement** |  |
| Resolution (Å) | 24.56 - 2.40 |
| No. reflections | 48,073 |
| *R*<sub>work</sub> / *R*<sub>free</sub> | 0.175/0.224 |
| No. atoms |  |
| Protein | 7,629 |
| Ligand/ion/solutes | 365 |
| Water | 275 |
| *B*-factors (Å<sup>2</sup>) |  |
| Protein | 32.99 |
| Ligand/ion/solutes | 47.01 |
| Water | 34.31 |
| R.m.s deviations |  |
| Bond lengths (Å) | 0.008 |
| Bond angles (°) | 1.021 |

**Table S2.** Transcript gene counts for BCO proteins in guts of wild type lancelet.

| gene name | Transcripts gene counts |  |  | Fixing acidic residues in active site |  |  |
|---|---|---|---|---|---|---|
|  | lancelet4 | lancelet 7 | Lancelet9 | **1** | **2** | **3** |
| LOC118409706 (BCOLg, XP_035666855.1) | 432 | 81 | 230 | **D** | **D** | **E** |
| LOC118412454 (BCOLb, XP_035671225.1 | 470 | 1761 | 304 | **D** | **D** | **E** |
| LOC118405420 (BCOLd) (XP_035660812.1) | 625 | 952 | 2049 | **D** | **E** | **E** |
| LOC118405416 (BCOLa1) (XP_035660808.1) | 147 | 386 | 401 | **D** | **E** | **E** |
| LOC118430142 (BCOLc, XP_035696745.1) | 115 | 154 | 66 | **D** | **D** | **E** |
| LOC118430143 (BCOLf (XP_035696746.1) | 47 | 78 | 86 | **D** | **E** | **E** |
| LOC118405340 (BCOa, XP_035660679.1) | 3557 | 13015 | 3106 | **E** | **E** | **E** |
| LOC118422787 (BCOb, XP_035686436.1) | 455 | 846 | 343 | **E** | **E** | **E** |
| LOC118426367 (BCOc, XP_035691598.1) | 581 | 2584 | 2067 | **E** | **E** | **E** |
| LOC118430335 (BCOLe, (XP_035697056.1) | 0 | 0 | 0 | **D** | **E** | **E** |

**Figure S1.**
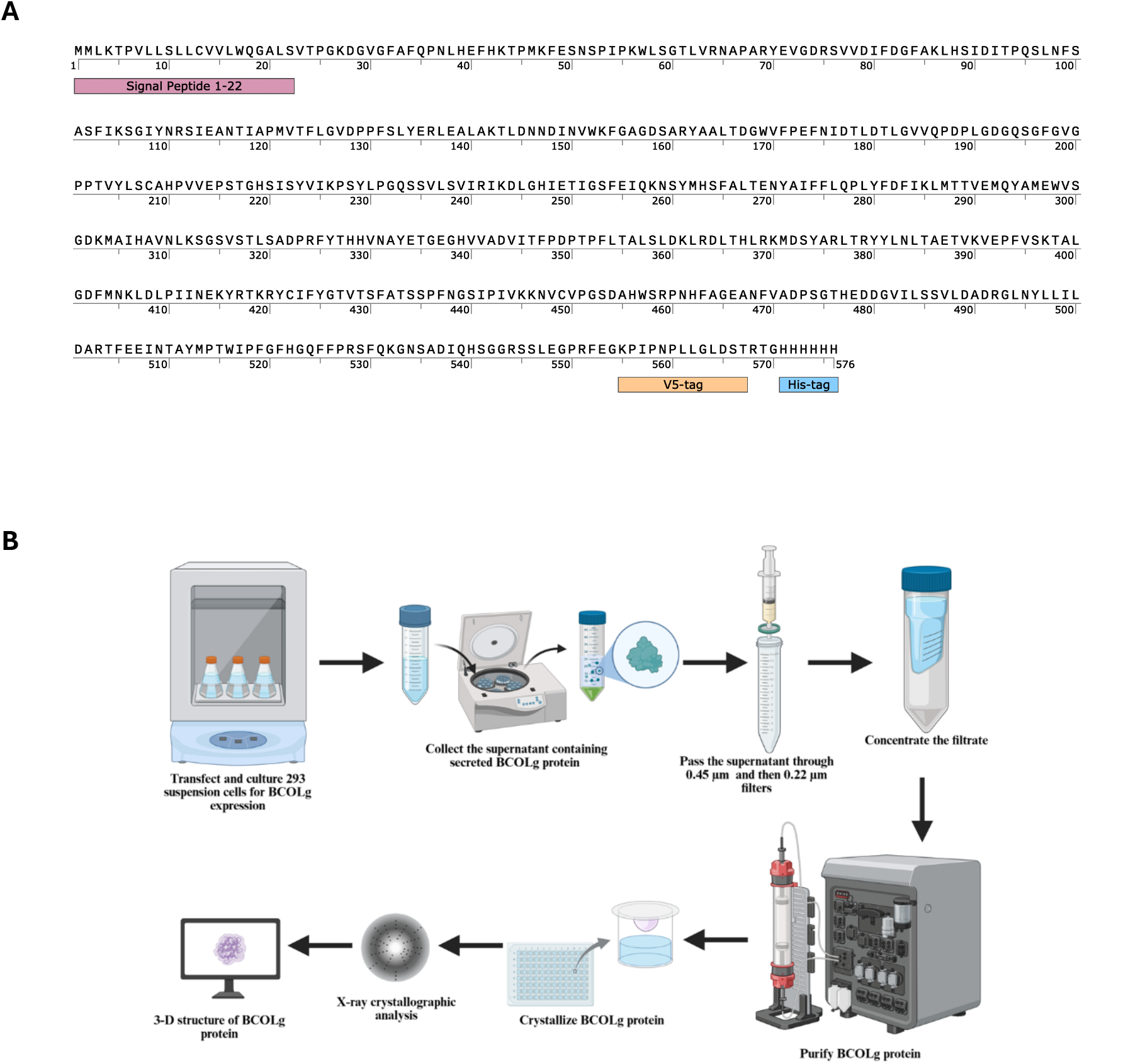
Production of recombinant BCOLg. (A) Schematic of the BCOLg construct cloned into the pcDNA3.1/V5-His-TOPO backbone, showing the signal peptide spanning the amino acid residues 1-22 and the C-terminal V5 and His tags. (B) Workflow for the expression and purification of recombinant BCOLg.

**Figure S2.**
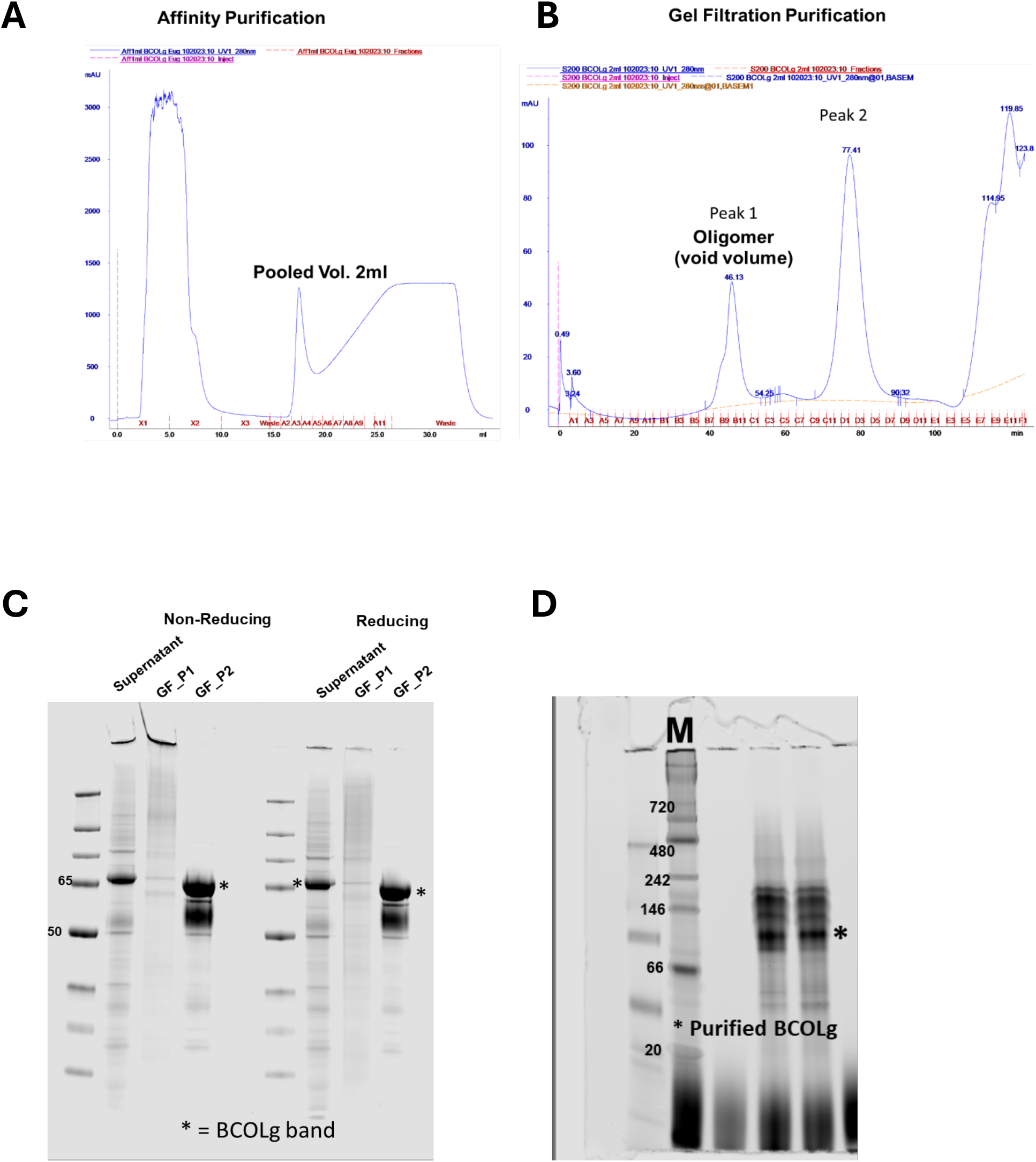
Production of recombinant BCOLg. (A) His-tag TALON (cobalt-based) affinity purification elution profile. (B) Subsequent gel filtration purification elution profile. (C) Noticeable protein degradation in peak 2 eluted from gel filtration column on SDS-PAGE reducing (left panel) and non-reducing (right panel) conditions. (D) Native gel for purified BCOLg showing the band corresponding to the dimer.

**Figure S3.**
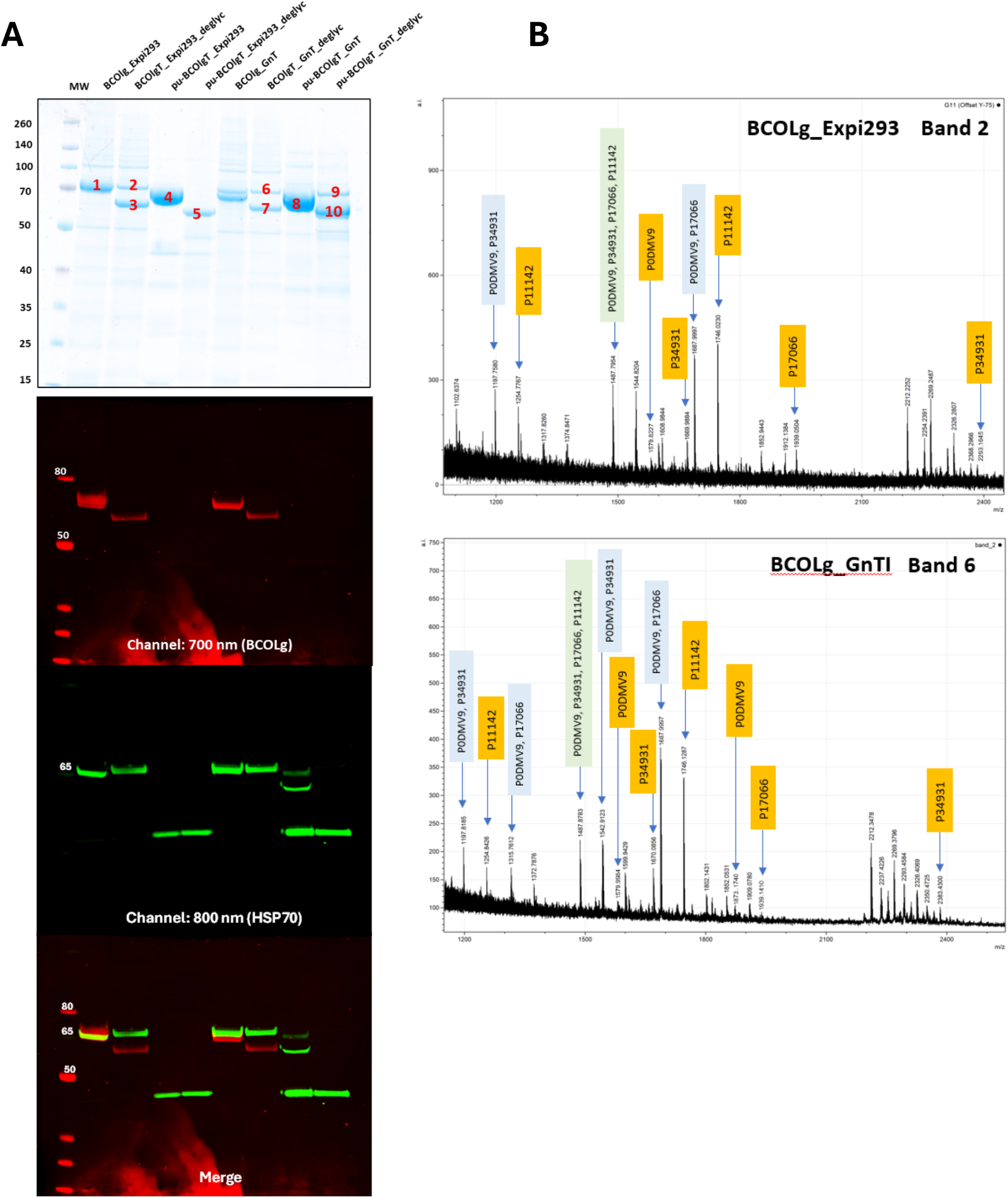
BCOLg is efficiently expressed and purified, and a distinct glycoform-dependent cell line is revealed by SDS-PAGE and mass spectrometry. (A) Coomassie-stained SDS-PAGE (upper panel) and western blotting (three lower panels) showing BCOLg and HSP70 in cell culture supernatants (Expi293 and GnTI-) and purified BCOLg samples before and after N-deglycosylation by PNGaseF treatment. Bands excised from the gel for additional analysis are numbered (1-10). Purified BCOLg was not identified due to the loss or masking of the C-terminal containing the His6 (Fig. S5) according to MS analysis (data not shown), and HSP70 was identified at a lower mass in comparison with the expected molecular weight, probably related to the degradation process according to results obtained for purified BCOLg from GnTI cells (pu-BCOLg_GnTI-). (B) MALDI mass spectrometry analysis for identification of contaminant HSP70 isoforms. Representative spectra from HSP70 in Expi293 (Band 2) and GnTI-(Band 6) samples show distinct peptide mass fingerprints, indicating different HSP70 isoforms, as annotated in each mass spectrum. Peptides assigned to HSP70 isoforms are labeled by their protein ID and color (yellow is for unique peptides for this isoform, blue is for peptides matching two proteins, and green for peptides for matching four proteins) PODMV corresponds to HSPA1B, P34931 corresponds to HSPA1L (confirmed by western blotting), P17066 corresponds to HSPA6, P11142 corresponds to HSPA8.

**Figure S4.**
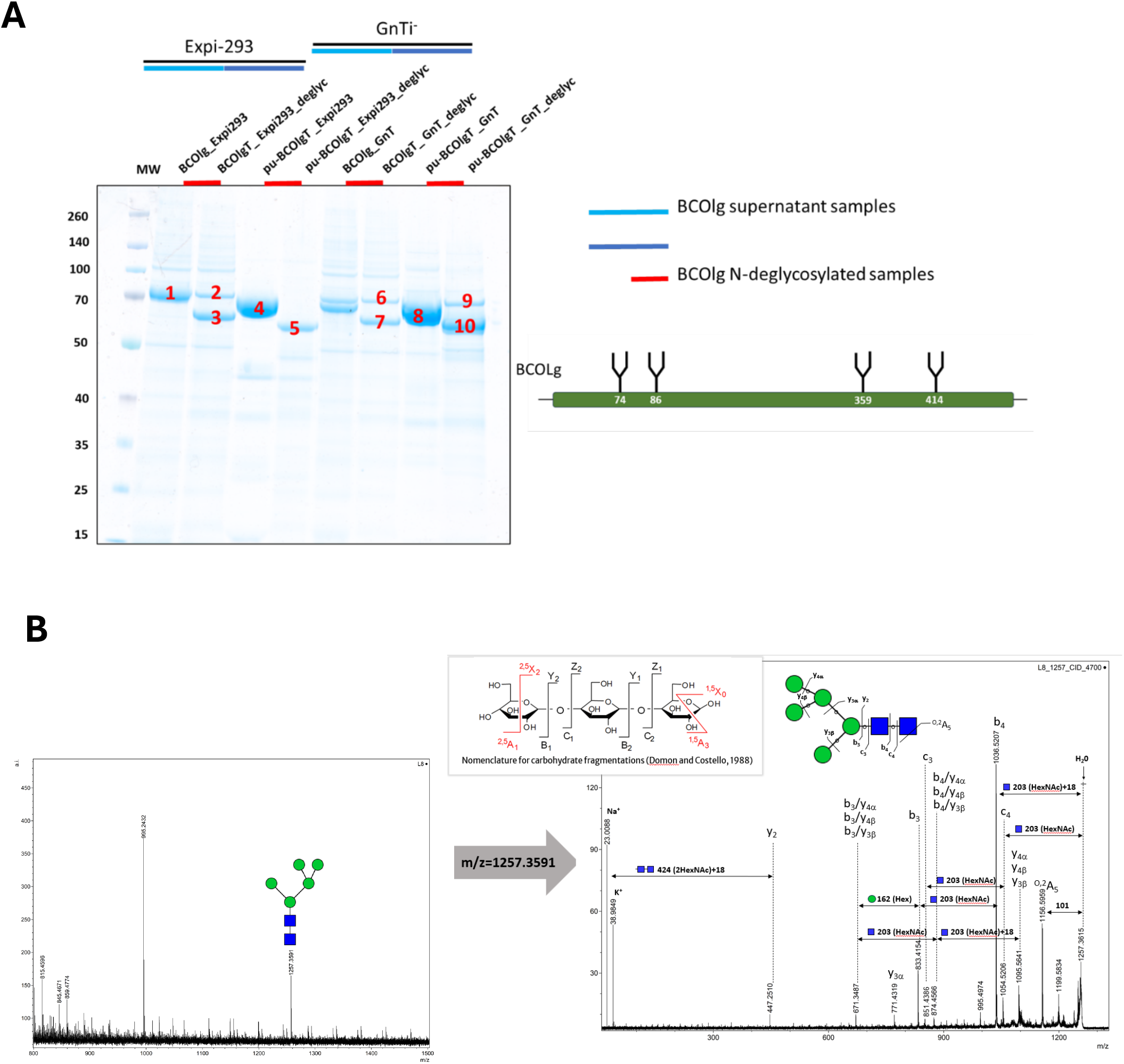
N-glycosylation analysis of BCOLg by gel electrophoresis and mass spectrometry. (A) Schematic representation of BCOLg indicating four N-linked glycosylation sites (N74, N86, N359, N414). (B) Coomassie-stained SDS-PAGE showing BCOLg in cell culture supernatants (Expi293 and GnT) and purified BCOLg samples before and after N-deglycosylation by PNGaseF treatment. Bands excised from the gel for additional analysis are numbered (1-10). (C) MALDI-MS spectrum of the released N-glycans from band 10. The two higher-intensity signals correspond to micro-cellulose (m/z 955.2432), used in the N-glycan purification protocol, and the oligomannosidic N-glycan structure (Man_5_GlcNAc_2_; m/z 1257.3591), which is the expected N-glycan structure linked to glycoproteins expressed in the GNTI cell line. (D) MALDI-MS/MS spectrum of the signal at m/z 1257.3591 in (C) to confirm the presence of Man_5_GlcNAc_2_ in BCOL expressed in the GNTI cell line.

**Figure S5.**
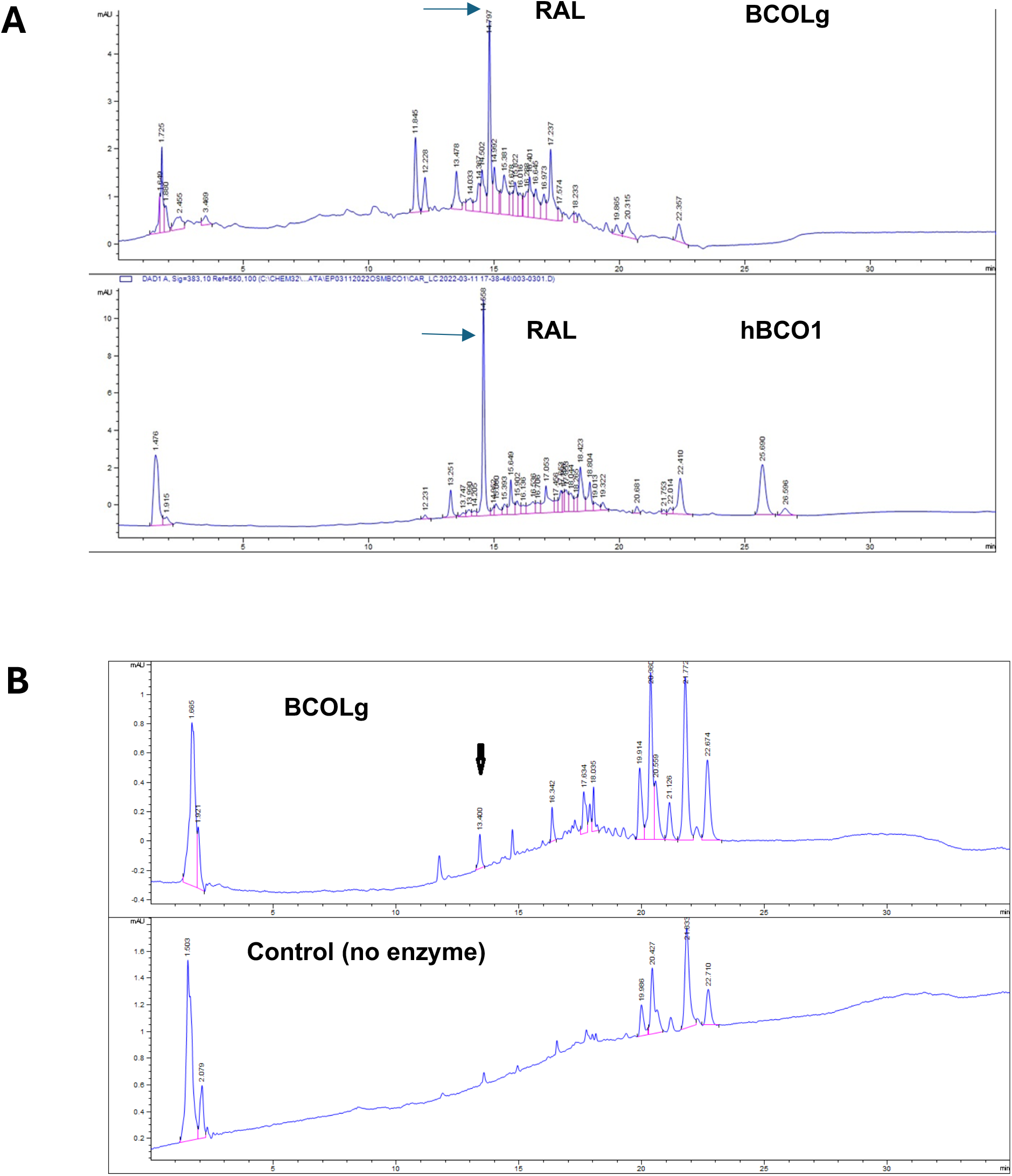
Comparison of BCOLg activity produced in Expi293 and GnTI-cells was compared to hBCOLg. (A) Comparison of BCOLg and hBCO1 activity in Expi293 cells, product of hBCO1 *al-trans* retinal is eluted at 14.6 min. (B) Activity of BCOLg produced in the glycosylation deficient cell line (GnTI-; upper panel) and no enzyme control (lower panel).

**Figure S6.**
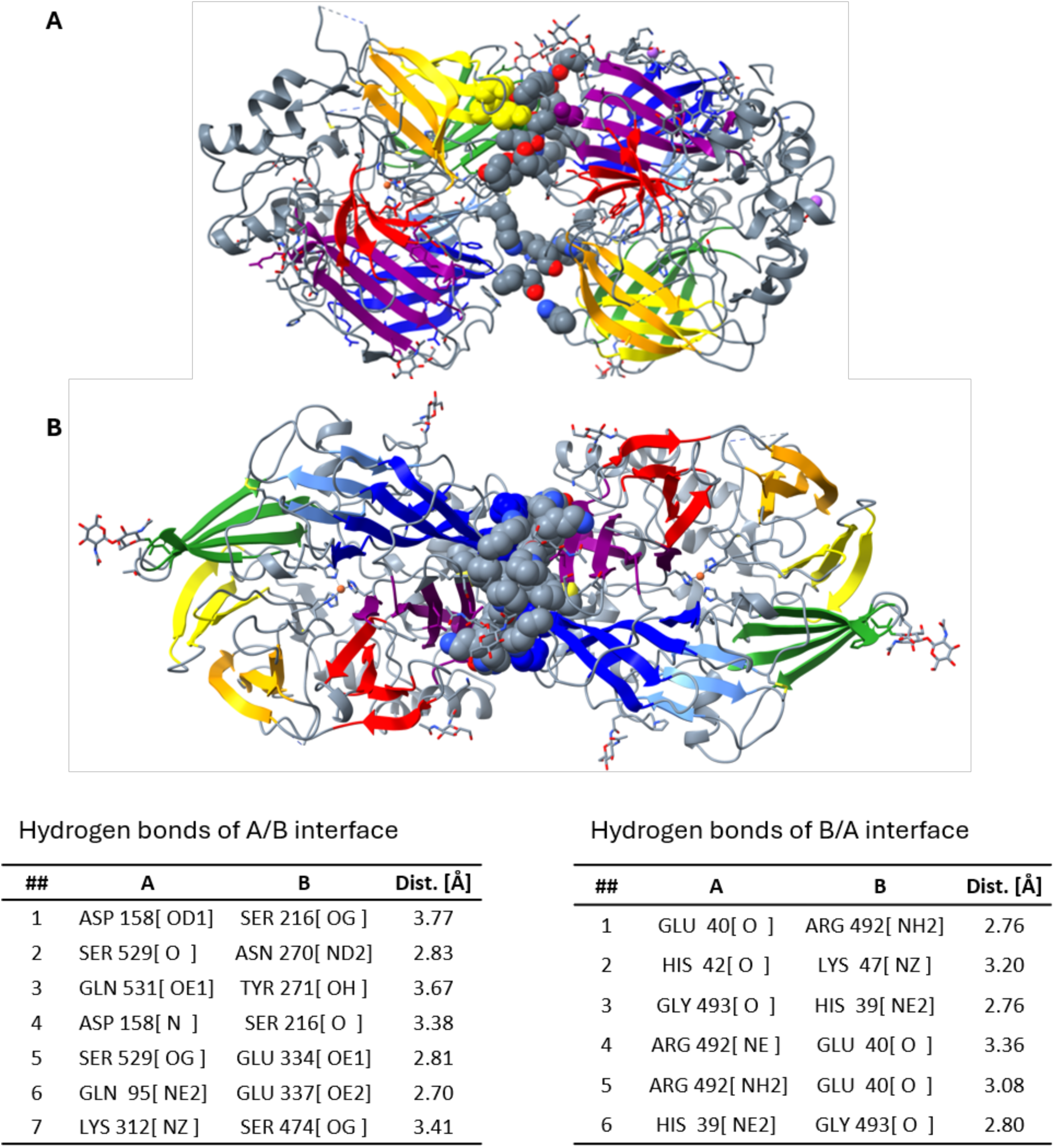
The two largest interfaces observed in the BCOLg crystal. Interface residues are shown as spheres, colored according to the blade of the seven-bladed β-propeller to which they belong. (A) The asymmetric unit contains two BCOLg molecules arranged base-to-base. The A/B interface comprises seven hydrogen bonds (listed in the table below) and buries a total surface area of 1210 Å² (600 Å² + 610 Å²). (B) The second-largest interface in the crystal is formed between molecule A and a symmetry-related molecule B (distinct from the one in the asymmetric unit) and corresponds to a side-to-side packing of two seven-bladed β-propellers that contact each other through blades VI (indigo) and VII (violet). This interface comprises six hydrogen bonds (listed in the table below) and buries a total surface area of 1480 Å² (740 Å² + 740 Å²). (C) Tables listing the hydrogen bonds at the A/B and B/A interfaces. PISA analysis was carried out with Protein interfaces, surfaces and assemblies’ service PISA at the European Bioinformatics Inst_6_it_0_ute as described in Methods.

**Figure S7.**
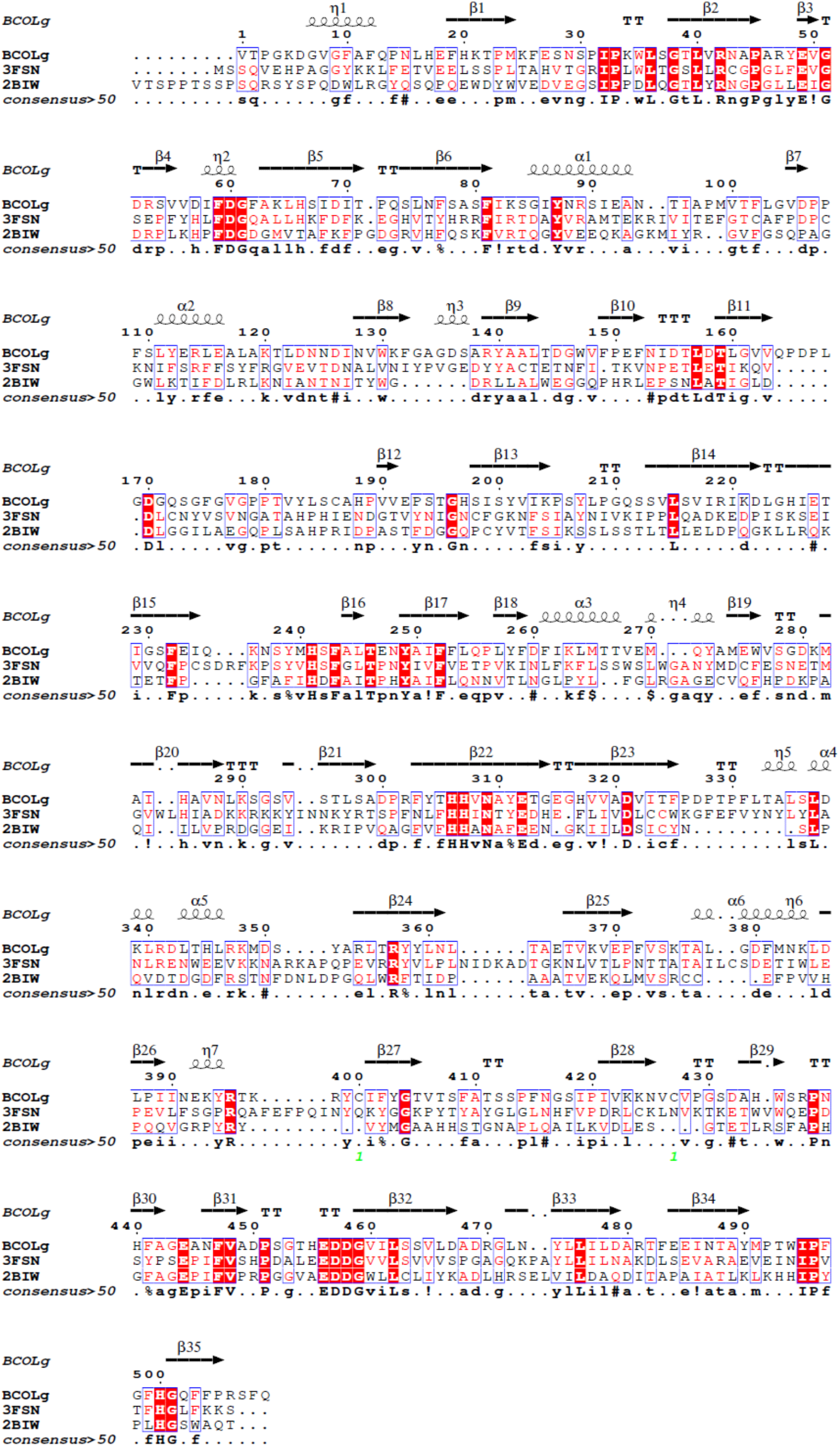
Sequence alignment of *Branchiostoma* BCOLg (PDB: 9PLV) with bovine RPE65 (PDB: 3FSN) and *Synechocystis* ACO (PDB:2BIW).

**Figure S8.**
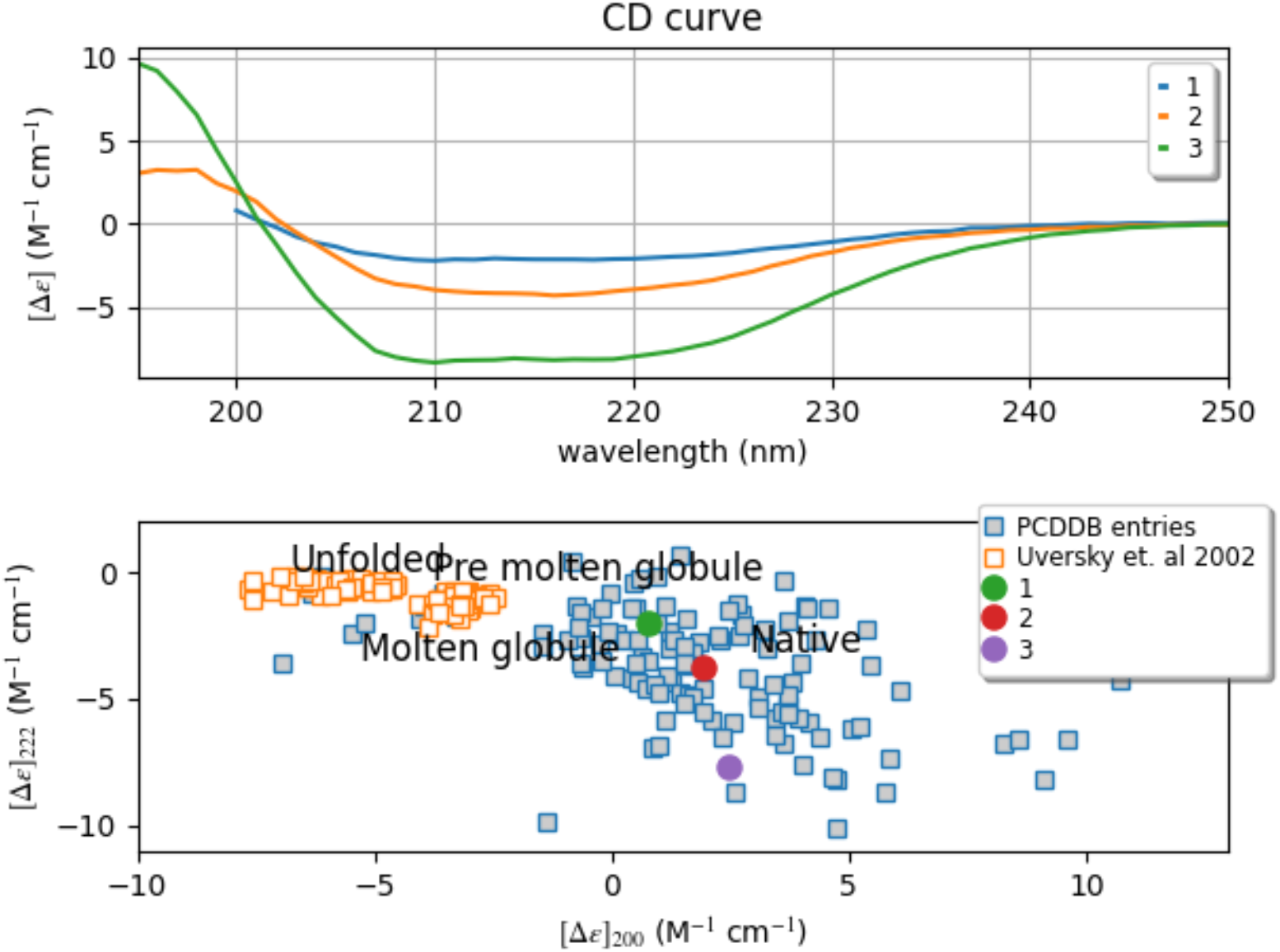
Far-UV circular dichroism analysis of BCOLg under different buffer conditions. Representative far-UV CD spectra (top panel) were collected between 195 and 250 nm for three BCOLg preparations. Protein 1 corresponds to BCOLg expressed in Expi293 cells and measured in Tris buffer containing glycerol. Protein 2 corresponds to the same Expi293-derived BCOLg measured in Tris buffer without glycerol following dilution. Protein 3 corresponds to BCOLg expressed in GnTI^−^ cells and measured in Tris buffer without glycerol. Both Expi293- and GnTI^−^-derived BCOLg proteins were purified by gel filtration in Tris buffer containing glycerol prior to CD analysis. The bottom panel shows the projection of the corresponding CD spectra onto the Protein Circular Dichroism Data Bank (PCDDB) reference map using [Δε]_222nm_ and [Δε]_200nm_ values. Blue squares represent reference proteins from the PCDDB dataset, defining regions associated with native, molten globule, pre-molten globule, and unfolded conformational states. All three BCOLg preparations cluster within the native region, indicating preservation of the overall β-sheet-rich β-propeller fold despite subtle differences in spectral intensity.

**Figure S9.**
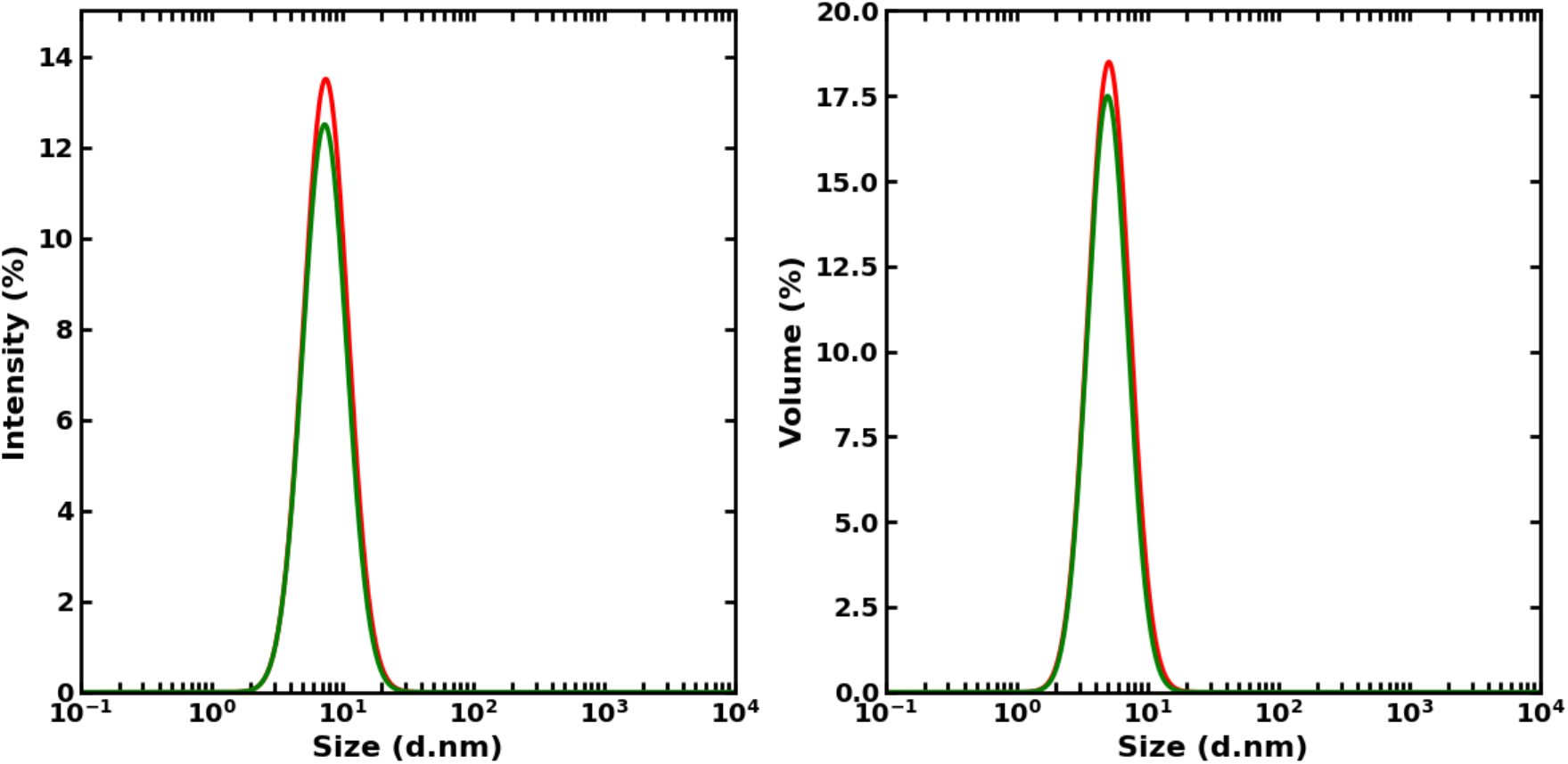
An example of dynamic light scattering measurement of BCOLg. BCOLg (6.8 mg/mL) used for crystallization was tested for homogeneity. PDI= 0.257, d= 6.2 ± 2.4 nm

**Figure S10.**
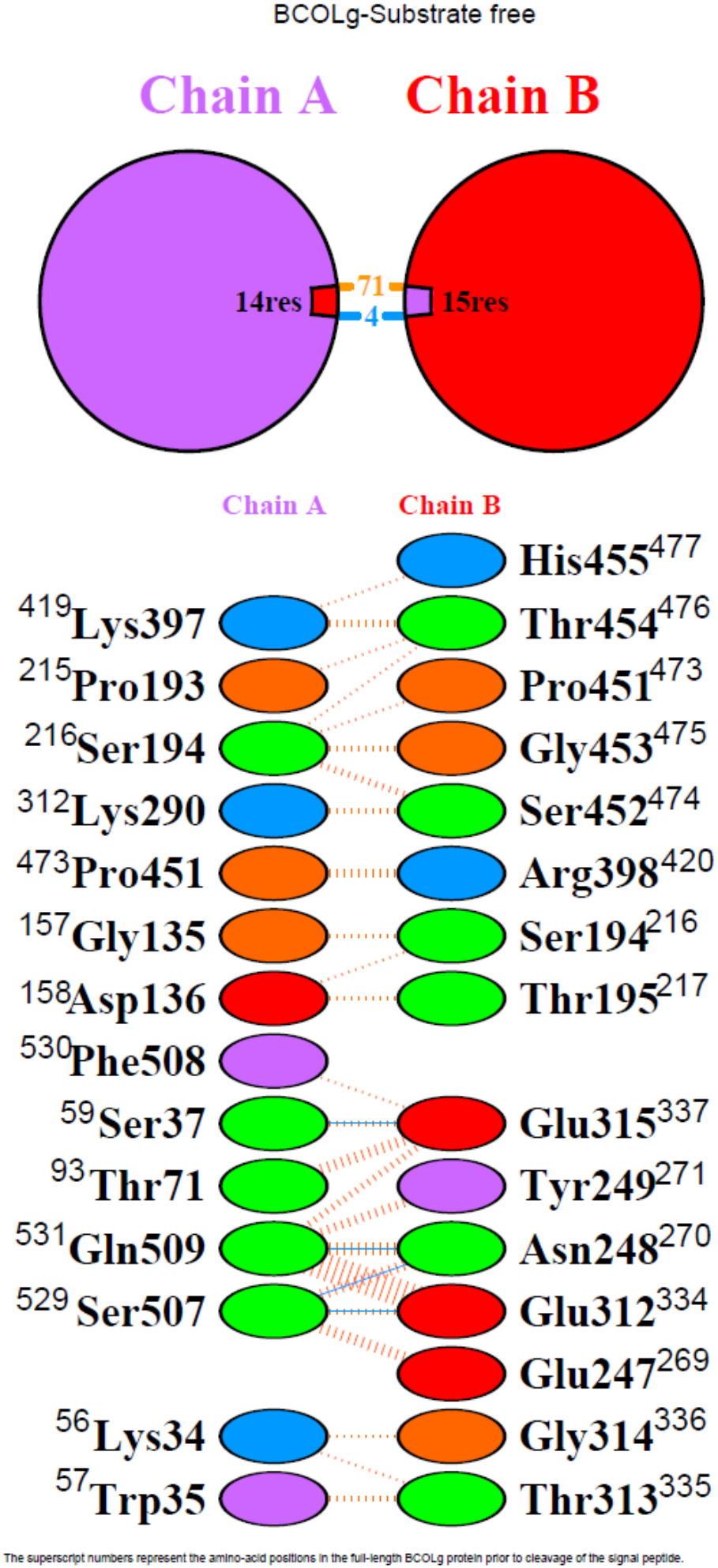
Interface analysis of dimeric BCOLg in the substrate-free crystal structure using PDBsum web server. Schematic representation of the interface between Chain A and Chain B in the substrate-free BCOLg crystal structure. The top panel summarizes the interface composition, showing that 14 residues from Chain A and 15 residues from Chain B contribute to the dimer interface, including multiple intermolecular contacts. The bottom panel details the specific residue–residue interactions across the interface, highlighting hydrogen bonds and other non-covalent interactions between symmetry-related chains. Residues are labeled with their positions in processed protein starting at position 23 as 1, and superscript numbers indicate their positions in the full-length BCOLg sequence prior to signal peptide cleavage.

